# FixGrower: An efficient and robust curriculum for shaping fixation behavior in rodents

**DOI:** 10.1101/2025.09.12.675850

**Authors:** Jessica R. Breda, Julie A. Charlton, Jovanna M. Willock, Charles D. Kopec, Carlos D. Brody

**Author notes:** equal contribution (Jessica R. Breda, Julie A. Charlton).

## Abstract

Center-port fixation is a common prerequisite for many freely-moving rodent tasks in neuroscience and psychology. However, typical protocols for shaping this behavior are non-standardized and inefficient. Moreover, motor errors in fixation termed ‘violations’ often account for a significant fraction of experimental trials, leading to notable data loss during experiments. In light of this, we developed FixGrower, a standardized protocol for center-port fixation training. FixGrower *(1)* requires a longer initial fixation requirement, *(2)* increases the required fixation duration at session boundaries customized to each animals’ performance, and *(3)* delays the introduction of violation penalties until the end of training. We demonstrate FixGrower decreases training time by 61%, yields low violation rates, and generalizes across rodent species and task difficulty. Moreover, the success of this curriculum is well supported by theories of operant conditioning and reinforcement learning. Our findings establish FixGrower as an efficient and broadly applicable curriculum for training fixation behavior in rodents, thereby accelerating training of many tasks in the field.

---

Training animals to perform perceptual decision making tasks in controlled laboratory environments has led to significant advancements in the fields of neuroscience and psychology (Oesch et al., 2024). Teaching animals these complex tasks takes a considerable amount of time, usually on the order of weeks to months (Kopec et al., 2024), and this has led to a body of theoretical and behavioral research focused on optimizing task learning.

These types of tasks typically have two components to learn: a perceptual stimulus rule and a corresponding motor sequence. The motor sequence is composed of actions related to both trial engagement and timing (e.g. withholding movement during the stimulus) as well as choice reporting (e.g. moving to the left or right given the stimulus). Most efforts to improve training efficiency have focused on accelerating stimulus rule acquisition by using a variety of techniques such as passive stimulus exposure (Schmid et al., 2024), or difficulty scheduling (Reuschenbach et al., 2023; Soma et al., 2014). In contrast, inefficiencies in motor sequence learning have been comparatively understudied, despite their considerable impact on overall training duration and performance quality.

In perceptual decision-making tasks, these structured, sequential motor behaviors are essential for ensuring consistent engagement and reliable measurement of stimulus-driven choices. In freely-moving versions of these tasks in rodents, a common motor requirement is “center-port fixation”. This is when the animal must insert and hold its nose in a central port (‘poke’) for a specified duration prior to and/or during stimulus presentation. Acquiring stable fixation behavior represents a surprisingly difficult challenge. For example, Schmid et al. (2024) reported that mice required 9–12 days of training to achieve a 250 ms pre-stimulus fixation hold. To learn a sound categorization rule following the hold took an additional 7–15 days. These findings underscore that the motor control demands associated with fixation can be substantial, sometimes requiring as much time as mastering the stimulus-response association itself.

Despite the widespread use of fixation in rodent decision-making tasks, training procedures for this motor requirement are often idiosyncratic across laboratories and minimally described in the literature (to highlight the variety in reporting and training across similar labs see Akrami et al., 2018; Constantinople et al., 2019; Jaramillo & Zador, 2014; Mah et al., 2023; Odoemene et al., 2018; Schmid et al., 2024; Scott et al., 2015) Nevertheless, several common elements are shared across approaches. First, the initial fixation requirement typically starts at an imperceptible threshold on the order of tens of milliseconds. Second, the required fixation duration increases gradually on a trial-by-trial basis. Third, failed fixation attempts are typically followed by either an inter-trial interval (ITI) delay or an additional penalty timeout that prevents an immediate retry after failure. Only after an animal consistently achieves the desired fixation duration are stimuli and the associated discrimination rules introduced.

However, even after animals consistently perform the stimulus discrimination rule, fixation errors, termed *violations* represent a persistent challenge. These events are common, accounting for approximately 20–35% of initiated trials (Ashwood et al., 2022; Constantinople et al., 2019; Cruz et al., 2022; Jang et al., 2024; Jaramillo & Zador, 2014; Licata et al., 2017). In other task settings, it has been shown that the ability to properly withhold movement is correlated with increased performance on the stimulus rule (Reuschenbach et al., 2023). In addition to their impact on behavior, violations also reduce data yield, as aborted trials are typically excluded from behavioral and neural analyses, thereby complicating dynamic models of decision making (Roy et al., 2021). Therefore, in the absence of standardized fixation training procedures, strategies for minimizing violations remain poorly defined.

In light of these challenges, we sought to develop an effective and efficient curriculum for shaping fixation behavior in freely moving rodents. Our goal was to design a robust, practical training procedure that could generalize across tasks and laboratories. We prioritized two outcomes: 1) minimize the time required to achieve a target fixation duration and 2) maintain low violation rates once at the target. There were three primary changes we implemented in this novel curriculum, which we refer to as “FixGrower”. First, the starting fixation requirement used on the initial day of training was considerably long at 400 ms. Second, we kept the fixation requirement stable within a session and grew only between sessions. The magnitude of the growth of the fixation requirement of each session was adaptively set by the performance of the animal in the preceding session. Finally, we implemented a time-out violation preceding an inter-trial-interval only after animals reached their target fixation requirement of 2 seconds, thereby allowing them to immediately retry following a failure during fixation growth.

Altogether, when compared to a “Legacy” curriculum used in our laboratory and similar to those in the field, we demonstrate this new approach is 61% faster, while maintaining low violation rates of 13.8% on average. Critically, we demonstrate our results are robust over time, species and even in later stages of task learning when a stimulus rule is introduced during fixation. We find the principles we implemented are well supported by theories in operant conditioning, reinforcement learning and curriculum learning. We believe these findings will expedite the necessary, foundational aspects of early task shaping and will be of use in a variety of rodent tasks.

## Results

### Fixation curricula comparison

We trained freely moving rats to perform a center-port fixation task (**Fig. 1a**), which is a common motor requirement of perceptual decision making tasks. In our task, animals poked and held their nose in an illuminated center-port until an auditory white noise go-cue indicated the end of the fixation period. If animals successfully fixated until the go-cue, they could collect water reward at a randomly illuminated lateral choice port. If animals violated the rule by failing to fixate until the go-cue, the trial was aborted. Depending on the curriculum and training stage, a 2-second timeout penalty and inter-trial interval delay may follow a violation. Required fixation durations ranged from 10 milliseconds to 2 seconds.

**Figure 1.**
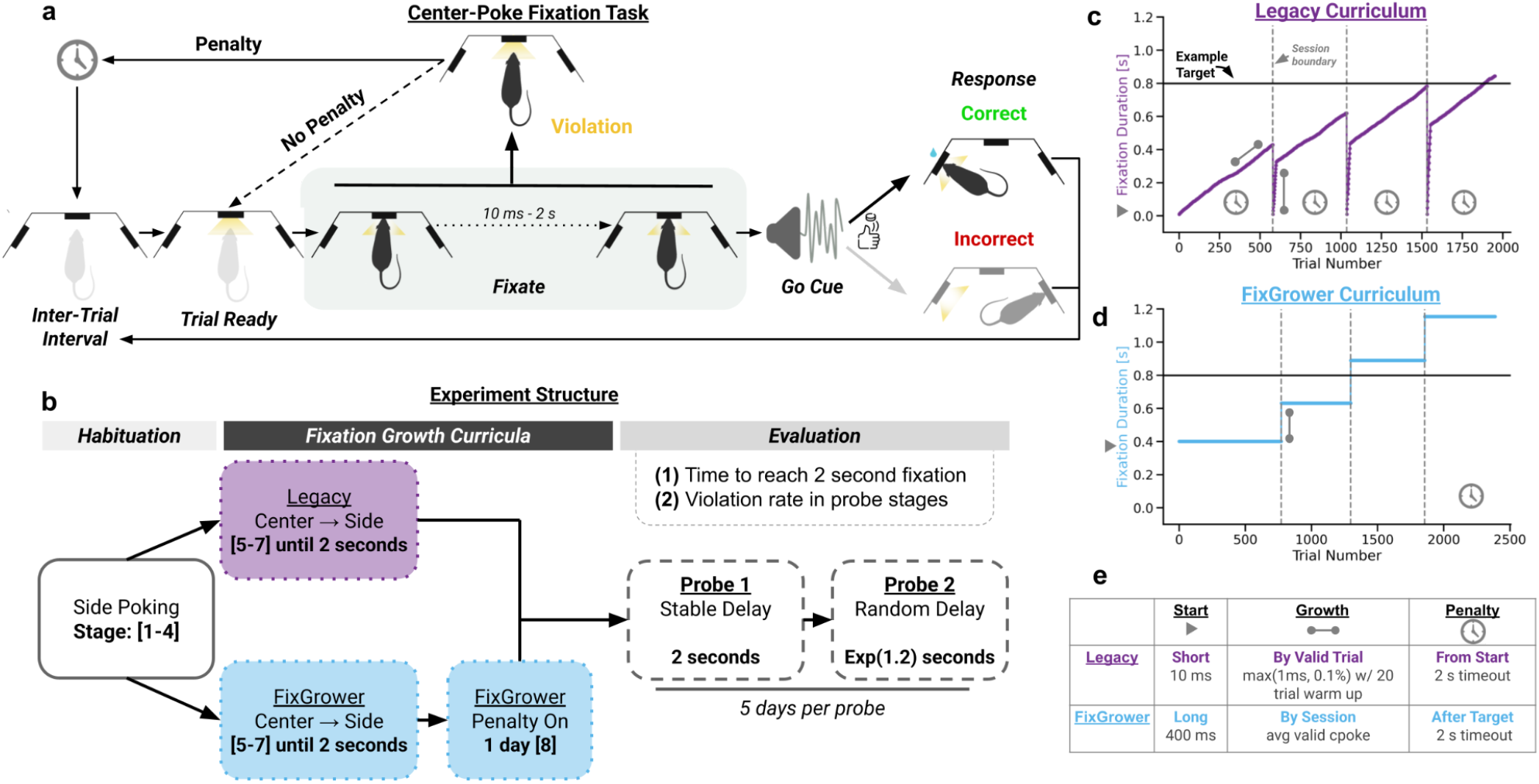
Overview of task structure and curricula. **(A)** Trial structure for center-port fixation task. Center light indicates trial ready, fixation occurs with nose in center hold and movement is allowed after a go-cue (200 ms white noise). The correct lateral choice is indicated by an illuminated side-port after the go-cue. If the animal prematurely removes their nose from center prior to the go-cue, the trial is considered a violation. A 2-second timeout penalty may follow a violation trial depending on the curriculum. An inter-trial interval period always follows a valid trial when animals fixate until the go-cue as well as trials when a violation penalty occurs. **(B)** Overview of curricula stages for our experiment. All animals have the same side poking habituation stages (gray bar) followed by differing fixation growth curricula (black bar) for Legacy (dark purple) and FixGrower (light blue) animals. Once the fixation target of 2 seconds is reached and violation penalty is turned on, all animals in the experiment go through 5 days of each probe stage to evaluate performance (gray bar). **(C)** Demonstration of how required fixation duration grows given the Legacy curriculum growth algorithm for four initial sessions (indicated by dashed vertical lines) with an example target duration of 0.8 seconds. **(D)** As in (C) but for the FixGrower curriculum growth algorithm. **(E)** Table summarizing differing features in the Legacy and FixGrower curricula.

We compared two different curricula for growing the fixation duration to a target hold of 2 seconds (**Fig. 1b**; **Table S1**). We evaluated the curricula using two primary metrics: (1) the number of days it took to reach the target and (2) violation rates in a series of two 5-day probe stages once the target was reached. The first probe stage tested performance at stable, long fixation holds of 2 seconds each trial. The second probe stage tested performance at variable fixation holds, sampled each trial from an exponential with a time constant (*τ*) of 1.2 seconds. Prior to the center-port fixation task, we trained all animals on a simpler side-poking task to habituate them to the behavior box environment and stimuli, and ensure they were motivated to collect water reward (**Table S2**; see *Methods: Curricula: Side Poking*). We then randomly assigned animals into one of two curricula: Legacy (*N = 8*), which is based on a typical fixation curriculum that is used in our laboratory and similar to those in the field (**Fig. 1c**), or FixGrower (*N = 9*),the novel fixation curriculum we present here (**Fig. 1d**).

There were three differences between the two curricula (**Fig. 1e**). First was the *starting fixation requirement*. This is the duration of time animals must fixate in the center-port prior to the go-cue on their first day of the curriculum. The Legacy curriculum had a “short” starting fixation requirement of 10 ms, while FixGrower had a “long” starting fixation requirement of 400 ms.

The second difference was the *fixation growth algorithm*. For Legacy, the required fixation duration grew following each non-violation (“valid”) trial by *max*(1 ms, 0.1%). Additionally, the required fixation duration started at 10 ms each session and “warmed up” over the first 20 trials to the maximum fixation duration achieved in the previous session. In contrast, FixGrower had a stable fixation requirement within a session that only grew between sessions to the average duration of an animal’s valid center-pokes in the previous session.

The final difference between the curricula was the *violation penalty introduction*. Legacy had an “early” penalty such that there was a 2-second timeout and inter-trial interval delay following a violation trial from the first day in the curriculum. In contrast, FixGrower had a “late” penalty such that there was no penalty for violations until the 2-second fixation target was reached. Therefore, prior to reaching the target, trials could be immediately restarted after a failed attempt. Once the target was reached, FixGrower had a single day of the violation penalty being turned on (Stage 8) before progressing animals into the probe stages. In contrast, the Legacy curriculum moved animals directly into the probe stages once they reached the target since the violation penalty was already on.

### FixGrower decreases training time

Since the center-poke fixation task is a motor prerequisite for many perceptual decision-making tasks, reducing the training duration for this phase is desirable to expedite progression to the main task of interest. Therefore, we assessed whether training speed differed between the two curricula. We found that FixGrower-trained animals grew significantly faster to the target fixation requirement with an average of 11.4 ± 5.4 days compared to Legacy-trained animals which averaged 29 ±18.3 days (*Mann–Whitney U* test, *U* = 65.5, *p* = 0.005; Post hoc power analysis; Cohen’s *d* = 1.33; power = 0.73; **Fig. 2a-b**). To accomplish this, FixGrower animals grew significantly more session-to-session than Legacy animals (mean ± SD: **L:** 0.07 ± 0.17 seconds/session, median : 0.07, *n*_Session_ = 245,vs. **FG:** 0.16 ± 0.17 seconds/session, median : 0.02, *n*_Session_ = 104; *Mann–Whitney U* test, *U =* 5654.0, p < 1.74 × 10^−12^; **Fig. 2c**).

**Figure 2.**
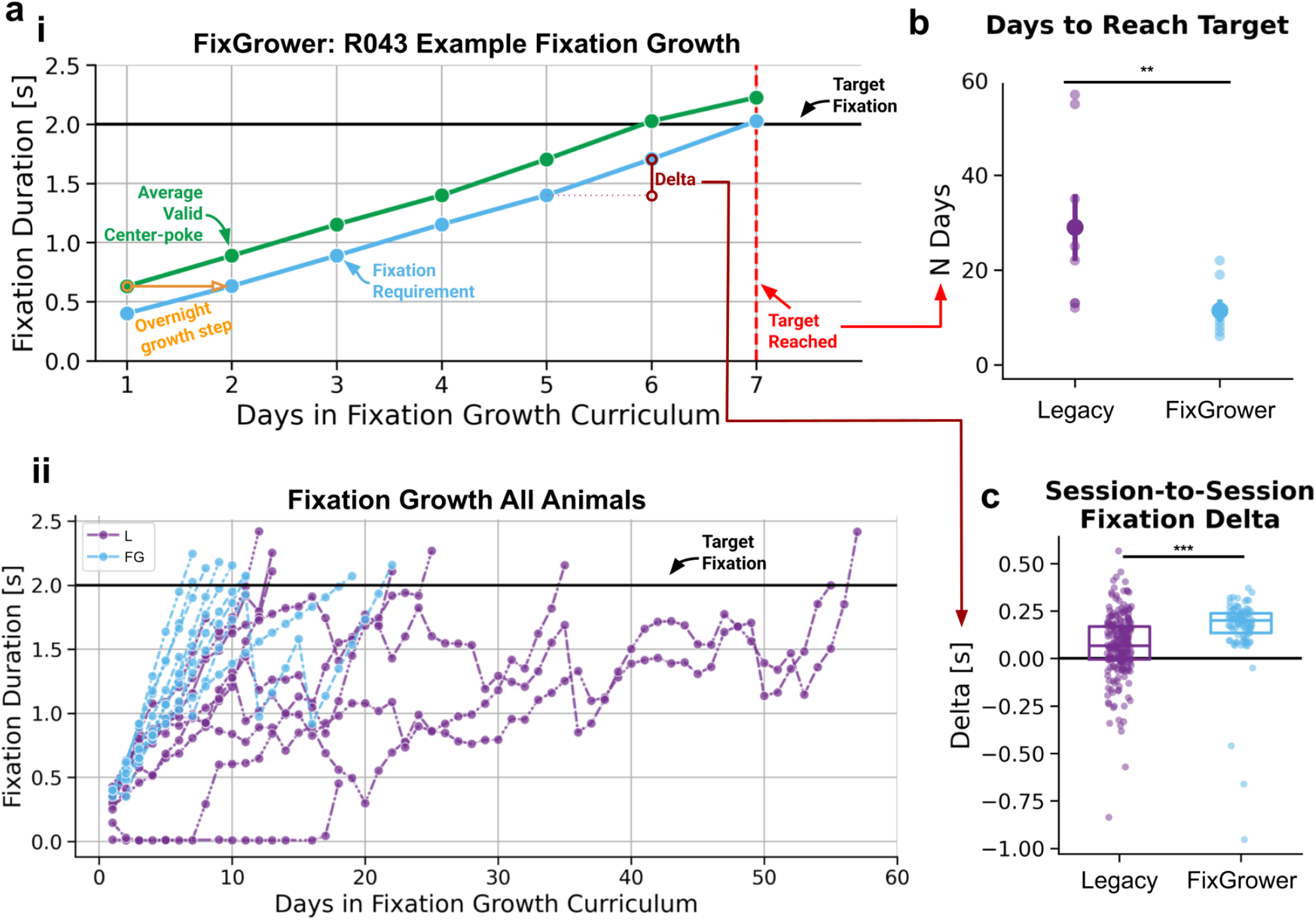
FixGrower-trained animals grow faster to fixation targets. **(A)(i)** Example growth trajectory for a single FixGrower animal. Black line indicates the 2-second fixation target. The light blue line indicates the required fixation for the given day. The green line indicates the average valid center-poke duration for a day, which is then set as the fixation requirement the following day (orange). **(A)(ii)** Maximum fixation duration reached in a session for each animal colored by experimental condition over days in the fixation growth curriculum. Black line indicates target fixation duration of 2 seconds. **(B)** Summary of days to reach 2-second target by fixation experiment for Legacy-trained animals (dark purple) and FixGrower-trained animals (light blue). Large circular points represent cohort mean and error bars represent standard error of the mean. Individual animals indicated by smaller points. Example days to reach the target indicated by the red dashed line in (A)(i). **(C)** Summary of fixation delta for each session by experimental condition. Boxes extend between lower and upper quartiles with a line at the median. Example delta indicated by a maroon line with open circles in (A)(i). *Statistics*: In (B) and (C) * *p < 0*.*05, ** p < 0*.*01, *** p < 0*.*001* and *ns* is *non-significant* tested using Mann–Whitney U tests. A mixed linear model with a random intercept for animal ID yielded the same results for (C).

This increased efficiency was not due to differences in engagement or performance between the cohorts prior to starting the fixation growth (**Fig. S1**). Nor was it due to an intrinsic growth ceiling in the Legacy growth algorithm given the number of trials performed (**Fig. S2**). In fact, simulations matched to rats’ trial rate with perfect performance demonstrated that Legacy animals could reach the fixation target significantly faster than FixGrower animals given the number of trials they were performing (5.8 ± 1.4 days, *Mann–Whitney U* test, *U* = 5.000, *p* = 0.003; **Fig. S2c**). The only notable difference between the two cohorts was that Legacy-trained animals had significantly more behavior box (“rig”) switches than FixGrower-trained animals (mean ± SD: **L:** 1.62 ± 0.52 switches, **FG:** 1.0 ± 0.0), which might destabilize performance as the animal adjusts to a new environment (**Fig. S1d**). However, this is anticipated given the significantly longer training times for the Legacy curriculum cohort. Further analysis showed no correlation between switches and performance (**Fig. S3**). Altogether, these data demonstrate that the structure of the FixGrower curriculum facilitated faster fixation growth and significantly reduced training times.

### FixGrower yields low and predictable violation rates

We next sought to evaluate fixation performance by assessing violation rates during a series of two probe stages administered after each animal was trained on their respective curricula to reach the 2-second fixation target. Each probe stage lasted five days. The first probe stage tested performance under stable fixation demands: each trial had a 2-second hold requirement. This served two purposes: first, to assess performance at a sustained, effortful fixation duration; and second, to evaluate how Legacy-trained animals performed under a consistent, trial-invariant rule. Both curriculum groups performed well, with average violation rates of 15.5 ± 5.3% (median : 16.2%) for Legacy-trained and 15.6 ± 8.2% (median :12.4%) for FixGrower-trained, and no significant difference in performance (*Mann–Whitney U* test, *U* =947.5, *p*_holm_ = 0.466; **Fig. 3a**).

**Figure 3.**
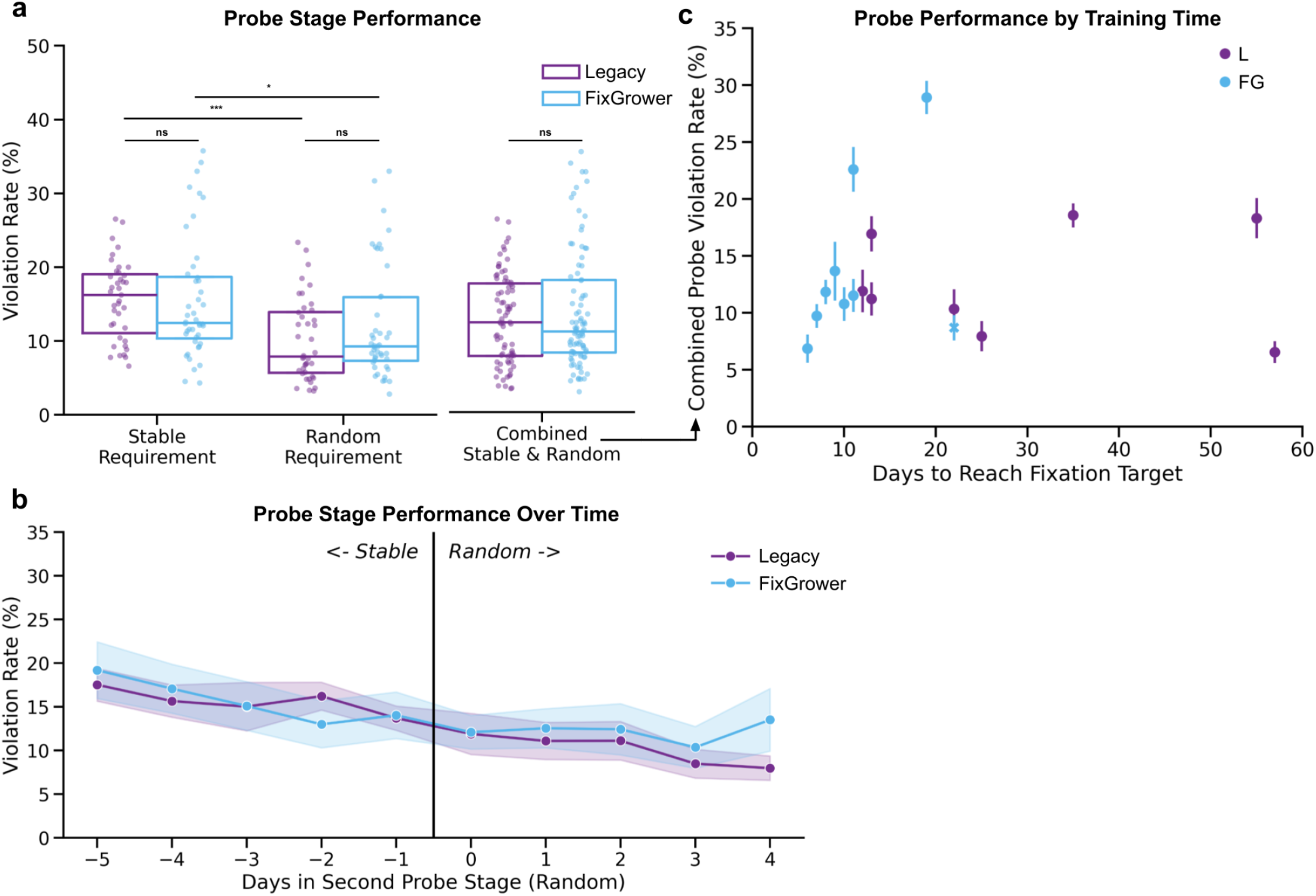
FixGrower-trained animals have low violation rates in probe stages that are predicted by training time. **(A)** Violation rate for the first, stable and second, random probe stages as well as the combined data from both stages for Legacy-trained (dark purple) and FixGrower-trained (light blue) animals. Dots represent individual animals and sessions. Boxes extend between lower and upper quartiles with a line at the median. Across-cohort statistics are computed using Mann–Whitney U test. Within-cohort statistics are computed using Wilcoxon signed-rank tests. Corrected for multiple comparisons with Holm method where * *p < 0*.*05, ** p < 0*.*01, *** p < 0*.*001* and *ns* is *non-significant*. **(B)** Violation rates over days in the probe stages for each cohort. Markers and lines represent the mean and shading indicates the standard error of the mean. For common alignment, day 0 is the first day of the random probe stage. **(C)** Relationship between days to reach target fixation of 2 seconds and average violation rate in the combined probe stages for each animal. Error bars are standard errors of the mean. A FixGrower outlier animal has an ‘X’ marker and is excluded from the mixed-effects linear model as discussed in *Methods: Excluded Animals*.

The second probe stage tested performance under variable fixation demands: on each trial, the required fixation duration was sampled from an exponential distribution with *τ* = 1.2 seconds, truncated to range between 1 and 2 seconds. This stage was designed to challenge FixGrower-trained animals, who had previously been exposed only to sessions with stable requirements. Nevertheless, they adapted well, showing a mean violation rate of 12.2 ± 7.7% (median : 9.29%) and no significant difference from Legacy-trained animals (mean ± SD: 10.1 ± 5.3%, median : 10.1%; *Mann–Whitney U* test, *U* =764.0, *p*_holm_ = 0.466; **Fig. 3a**). Further, both cohorts showed a downward trend in violation rates over the ten total probe sessions (**Fig. 3b**), and both exhibited significantly lower violation rates in the second, random probe stage compared to the first, stable one (*Wilcoxon signed-rank* tests, **L:** *W*= 1.0, *p*_holm_ = 0.046; **FG:** *W*= 2.0, *p*_holm_ = 0.046; **Fig. 3a**). These data demonstrate that the FixGrower-trained animals not only generalized well to variable fixation demands but also improved with continued exposure, indicating ongoing refinement of motor control even after reaching the training criterion.

Lastly, we were interested in the relationship between training times and performance, as it is of significant cost for experimenters to wait with high uncertainty about an animal’s training potential. To test this, we examined whether the number of sessions it took to grow to the target fixation could predict an animal’s combined probe-stage performance. While no relationship was observed in Legacy-trained animals, we found that FixGrower-trained animals who reached the target faster had lower violation rates in the probe stages (**Fig. 3c**). A mixed-effects linear model comparing probe-stage violation rate and the number of days to reach the target fixation requirement revealed a significant difference in slope between the cohorts (interaction term *β* = 0.016, *p* < 0.001, *Methods: Excluded Animals)*. These findings suggest that, in the FixGrower curriculum, longer training durations predicted poorer probe-stage performance, highlighting an exploitable trade-off between training speed and behavioral stability in this curriculum.

### FixGrower generalizes over time, tasks and species

Given the demonstrated success of the FixGrower curriculum, we next sought to test the robustness and generalizability of the FixGrower curriculum to determine its suitability for broader adoption in the field. We first assessed long-term task performance. Specifically, we examined whether animals trained on the FixGrower curriculum could sustain low violation rates over an extended period without further experimenter intervention. We selected a subset of animals trained on the FixGrower curriculum from the preceding fixation experiment (*N = 8*) and monitored their performance for 100 days following completion of the probe stages. Each day, the required fixation duration varied randomly for each animal and was sampled from an exponential distribution (*τ* = 1.2 seconds), truncated to range between 1 and 2 seconds (**Fig. 4a-b**). Across this prolonged evaluation, animals consistently maintained low and stable violation rates, averaging 9.89 ± 8.26 % (median7.27%; *n*_Session_ = 800; **Fig. 4c-d**). These findings strongly indicate that animals trained with FixGrower reliably mastered and maintained fixation task performance.

**Figure 4.**
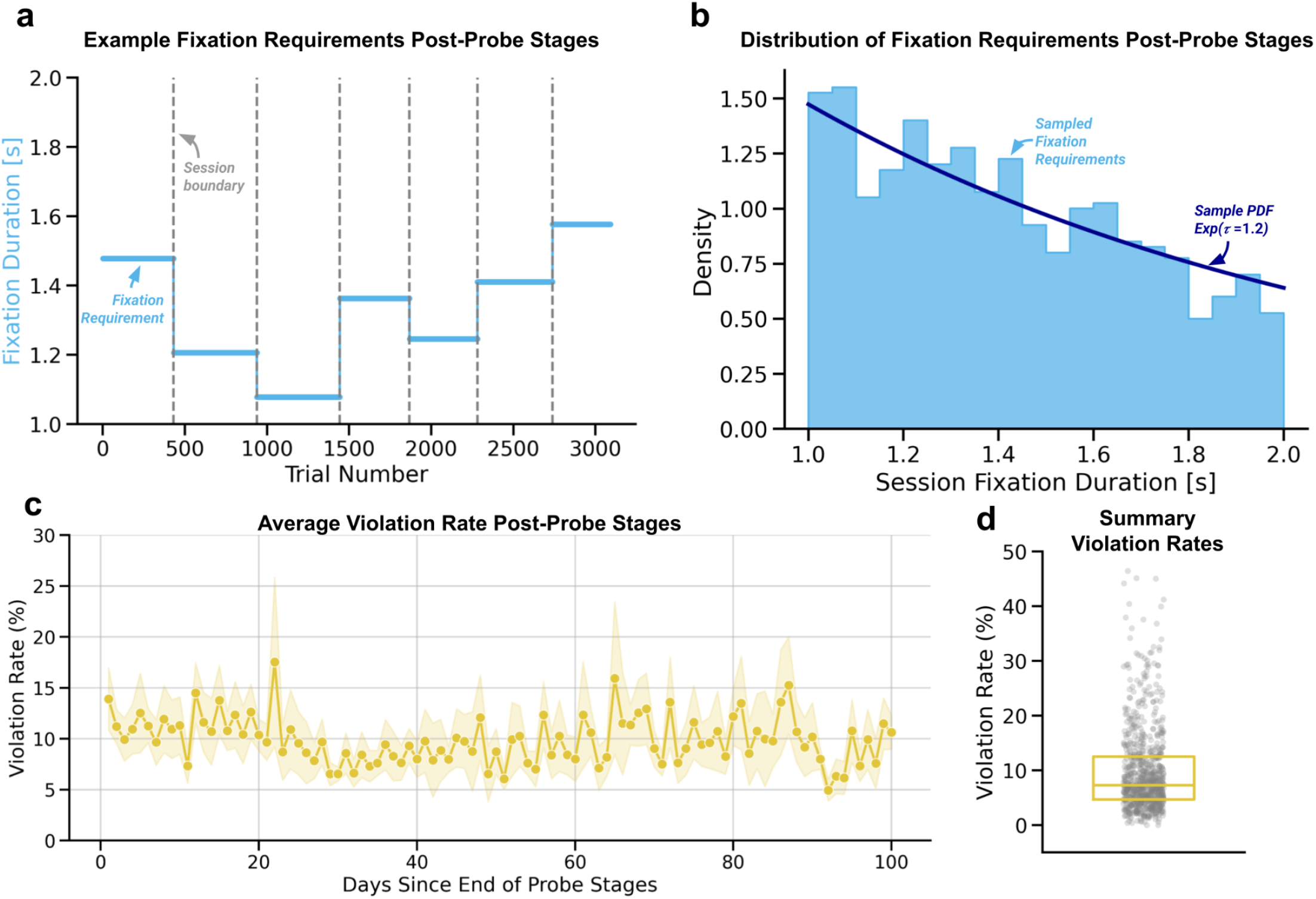
FixGrower-trained animals maintain robust task performance over extended periods. **(A)** Example sequence of fixation duration requirements sampled across trials from one representative animal (R053) after probe stage completion. Fixation durations were sampled from an exponential distribution (*τ* = 1.2 seconds) truncated between 1 and 2 seconds, updated each session. Dashed vertical gray lines indicate session boundaries. **(B)** Distribution of fixation duration requirements aggregated across all sessions. Histogram shows empirically sampled fixation durations (light blue bars), overlaid with the theoretical exponential PDF seconds) that fixation durations were drawn from. **(C)** Session average violation rates (%) plotted as a function of days post-probe stages completion. Each point reflects daily mean violation rates averaged across animals, with shaded regions indicating standard error of the mean (SEM). **(D)** Summary plot displaying individual session violation rates aggregated over the entire 100-day assessment period. Box extends between lower and upper quartiles with a line at the median and each gray dot represents a single session for a single animal.

Next, to test the generalizability of FixGrower, we evaluated whether its low violation rates would persist when animals had to fixate while processing task-relevant stimuli. We trained a separate cohort of rats (*N = 13*) on a sound discrimination task following FixGrower (see *Methods: Curricula: Sound Discrimination Task*). In this task, two 400 ms tones were separated by a 150 ms delay and flanked by additional delays (250 ms before and 450 ms after) for a total of 1.65 seconds of fixation (**Fig. 5a**). The first tone (3 kHz) served as a distractor, while the second tone indicated the rewarded side (12 kHz = right; 3 kHz = left; **Fig. 5b**) and a 200 ms white noise go-cue indicated the end of the fixation period. Using FixGrower, rats averaged 6.6 ± 2.2 days to grow to a common 1.45 s fixation target (min : 5, median : 6, max : 12; **Fig. 5c**). Once the sound rule was introduced, rats performed well in this task, achieving an average of 87.6 ± 5.9 % correct (*n*_Session_ = 170; *n*_trials_ = 39,028; **Fig. 5d**). Crucially, their violation rates remained low at 11.2 ± 9.5% (median : 8.33%; **Fig. 5e**), indicating that the FixGrower curriculum supports stable fixation even when the stimulus appears during the fixation period.

**Figure 5.**
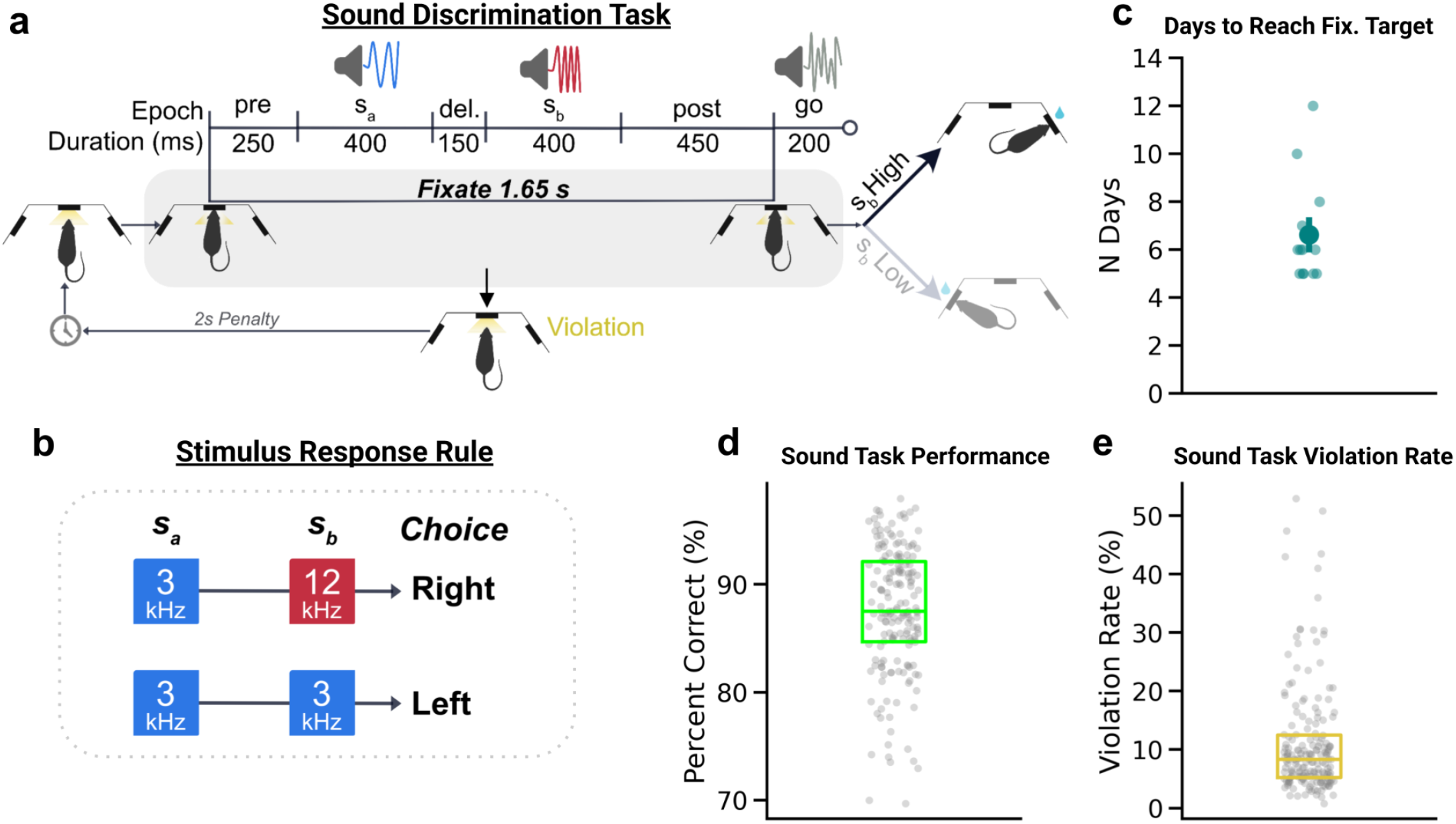
FixGrower supports efficient and stable fixation in a sound discrimination task. **(A)** Schematic of the sound discrimination task performed by a cohort of rats (*N = 12*) after completing FixGrower. Animals fixated for 1.65 seconds while auditory stimuli were presented: a non-informative distractor tone (s_a_) followed by a second tone (s_b_) indicating the correct choice (right for high frequency, 12 kHz; left for low frequency, 3 kHz). A 200 ms white noise go-cue signaled the end of fixation. **(B)** Stimulus set and response mapping used in the task. The frequency of s_b_ determined the rewarded port; s_a_ was always 3 kHz and non-informative. **(C)** Days required to reach the shared fixation target of 1.45 seconds (see *Methods: Curricula: Sound Discrimination Task*). Dots represent individual animals; the large dot shows the mean and error bars denote the standard error of the mean.**(D)** Task performance measured as percent correct on non-violation trials. Gray dots are individual animals and sessions; the green box shows the inter-quartile range with the median. **(E)** Violation rate for the sessions shown in (D). Gray dots are individual sessions; the yellow box shows the inter-quartile range with the median.

Finally, we evaluated whether the FixGrower generalizes effectively from rats to mice, as previous literature has demonstrated that mice typically require longer durations to learn fixation tasks and exhibit higher average violation rates (Jaramillo & Zador, 2014; Odoemene et al., 2018; Schmid et al., 2024). We trained a cohort of mice (*N = 5*) to fixate to a target duration of 1.1 seconds (**Fig. 6a**). We chose this target because it is a substantial hold length that has been previously used in the field with reported training times (Odoemene et al., 2018). On average, these mice reached the 1.1-second fixation target in 4.4 ±0.89 days (range: min =3, median = 5, max = 5 days; (**Fig. 6b**). Notably, a subset of these animals continued to grow beyond the initial target, achieving fixation durations exceeding 2 seconds within 9 days (**Fig. S4a**). Additionally, FixGrower-trained mice exhibited session-to-session fixation growth rates (0.25 ± 0.08 seconds/session, median : 0.21, *n*_Session_ = 22; **Fig. 6c**) similar to those observed in FixGrower-trained rats (0.16 ± 0.17 seconds/session, median : 0.20, *n*_Session_ = 104). These results reflect substantial improvement over previously reported training times of approximately 10–12 days using a curriculum similar to Legacy (Odoemene et al., 2018).

**Figure 6.**
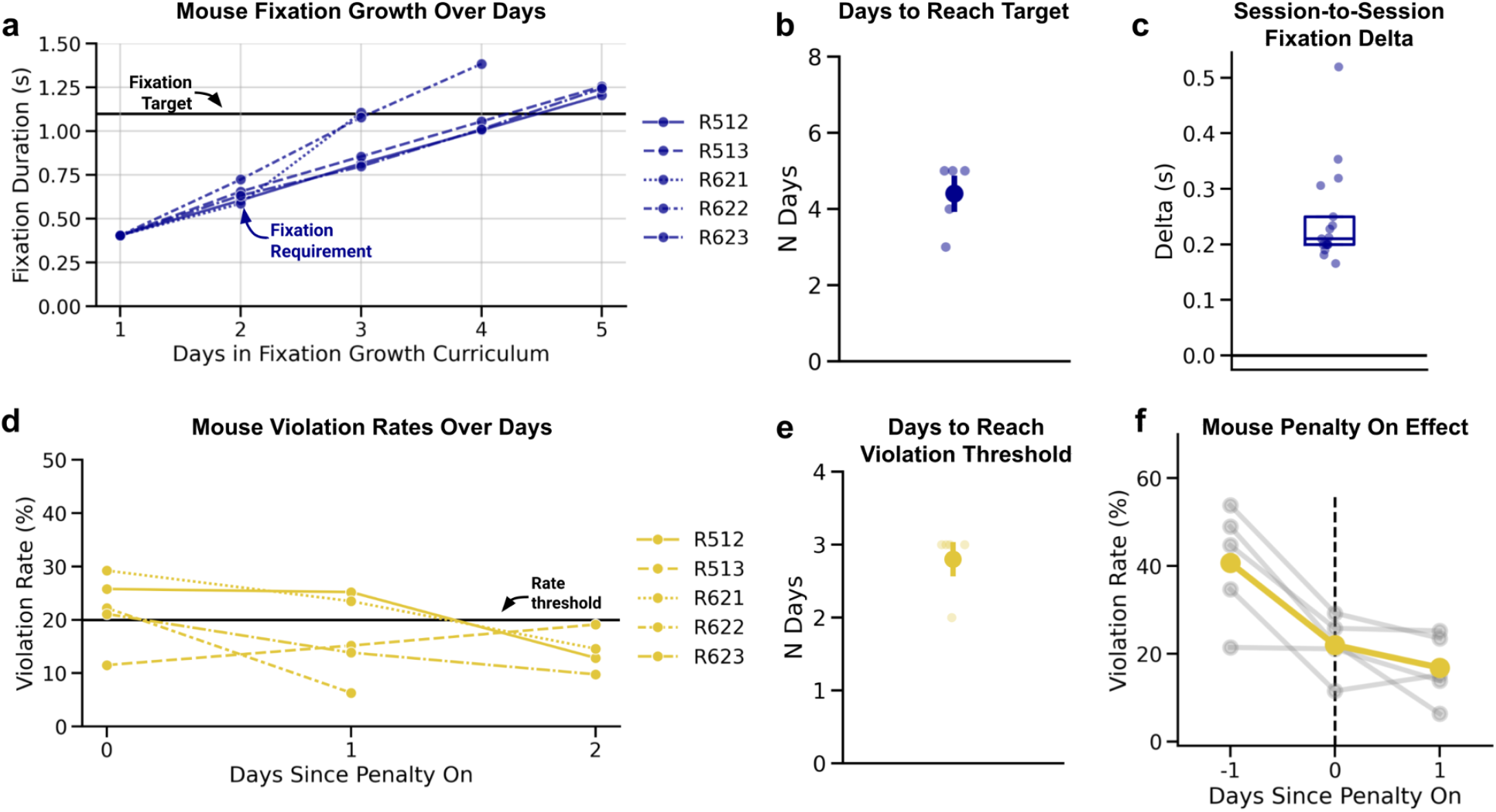
FixGrower Curriculum generalizes to mice. **(A)** Growth of session-maximum fixation duration across training days for individual mice (*N = 5*) using FixGrower. Each line represents one mouse. All animals were trained to a target of 1.1 seconds (black horizontal line). **(B)** Number of days each mouse required to reach the 1.1-second fixation target. Individual points represent mice; the large point indicates the cohort mean ± standard error of the mean (SEM). **(C)** Distribution of session-to-session fixation duration increases (delta), showing median (line), inter-quartile range (box), and individual animal and session values (dots). See **Fig. S4a** for delta intuition. **(D)** Violation rates for individual mice across days after introducing a 100 ms penalty. Each line represents one mouse, with a performance threshold of 20% indicated (black line). Mice complete this stage once they perform greater than 150 trials below this threshold. **(E)** Days required for each animal to achieve violation rates below 20% while maintaining 150 trials per session. **(F)** Behavioral response to penalty onset. Violation rates are shown on the day before (–1), the day of (0), and the day after (+1) penalty activation. Gray lines represent individual mice, and the bold yellow line shows the group mean.

We next assessed violation rates under modified conditions, using a short violation penalty (100 ms) and fixation durations ranging from approximately 1.5 to 2.7 seconds (see *Methods: Curricula: Mouse Fixation Experiment*, **Fig. S4b**). We determined how many sessions were required for animals to achieve violation rates below 20% while completing at least 150 trials per session (**Fig. 6d**). This 20% threshold represents the lower bound of violation rates commonly reported in prior mouse studies (Ashwood et al., 2022; Cruz et al., 2022; Jaramillo & Zador, 2014; Odoemene et al., 2018). Remarkably, all mice were sensitive to the short penalty and met this performance criterion within just 2–3 days (**Fig. 6e-f**), underscoring the effectiveness of the FixGrower curriculum in rapidly achieving low violation rates that match or exceed established benchmarks. Collectively, these findings indicate that FixGrower efficiently promotes rapid and robust fixation behavior in a variety of settings.

### Curriculum features modulate vigor, engagement stability, and motor control during learning

Having demonstrated that the FixGrower curriculum is more efficient than the Legacy curriculum and generalizes across diverse settings, we next sought to understand the factors underlying this success. In our final set of analyses, we examined how specific curriculum features—*starting fixation duration, fixation growth algorithm*, and *violation penalty introduction*—may have influenced behavior to either accelerate training in FixGrower-trained animals or impede it in Legacy-trained animals.

First, we focused on the delayed introduction of the violation penalty in FixGrower and its influence on motivation and trial initiation vigor. During fixation growth, the cost of failure is lower in FixGrower compared to in Legacy; in FixGrower, animals can immediately retry post-violation without additional delays, whereas in Legacy, animals incur both a 2-second violation penalty and an inter-trial interval delay. Notably, because fixation growth occurred on a per-trial basis in the Legacy curriculum, growth for these animals is constrained by the number of trials completed per session. We therefore hypothesized that the high cost of failure in the Legacy curriculum would suppress trial initiation, leading to fewer completed trials per session and thereby slowing the rate of fixation growth.

Consistent with this hypothesis, Legacy-trained animals indeed attempted significantly fewer trials per session compared to FixGrower-trained animals (**L:** mean ± SD 332 ± 109 vs. **FG:** 470 ± 184; *U* = 5689, *p* = 2.96 × 10^−17^; **Fig. 7a**), despite achieving a similar number of rewarded trials (**L:** 247 ± 80 vs. **FG:** 252 ±110; *U* = 12258.5, *p* =0.372; **Fig. 7b**). On average, FixGrower-trained animals reinitiated trials 3.3x faster after a violation than the Legacy-trained animals, even after accounting for experimenter-imposed delays (**L:** median3.82s, mean ± SD 16.6 ± 57.3s, *n*_trails_ = 17.908 vs. **FG:** median0.15s, mean ± SD 5.00 ± 53.0s, *n*_trails_ = 22,622; Mann–Whitney U = 3.21 × 10^8^, *p* < 0.001; **Fig. 7c**). This quick reinitiation time likely drove the higher trial initiation frequency observed in the FixGrower-trained animals.

**Figure 7.**
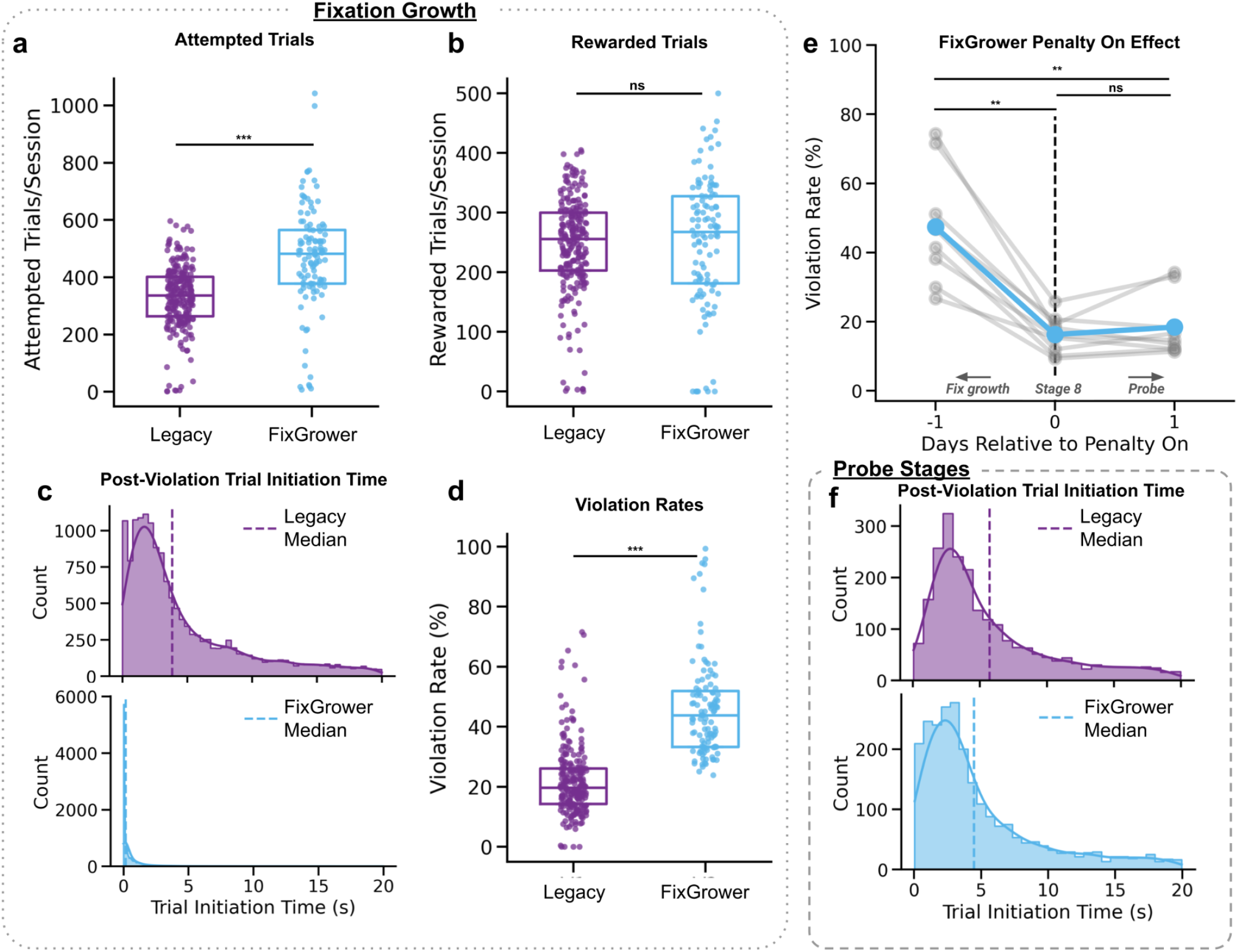
FixGrower-trained animals have more vigor than Legacy-trained animals. **(A)** Number of attempted trials per session for each cohort (Legacy: dark purple, FixGrower: light blue) during fixation growth where attempted trials are either violation or valid trials. **(B)** Same as **(A)** but number of rewarded trials per session where rewarded trials are valid hit trials. See **Fig. S1b** for hit rates. **(C)** Histogram and kernel density estimate of trial initiation times post-violation trials during fixation growth for each experimental cohort (Legacy: upper, FixGrower: lower) truncated to show initiations less than 20 seconds long. Note trial initiation times do not include incurred violation timeout penalty or inter-trial delay for Legacy-trained animals. Dashed lines indicate the cohort median. **(D)** Session violation rates during fixation growth for each experimental condition. **(E)** Violation rates for FixGrower-trained animals on the day before (−1), of (0), and after (+1) the 2-second violation penalty being turned on. Gray lines indicate individual animals. The light blue line indicates the group mean. Error bars indicate standard error of the mean. **(F)** As in **(C)** but with trials from the probe stages. *Statistics:* In **(A, B, D)** boxes extend between lower and upper quartiles with a line at the median and each dot represents a single session for a single animal. * *p < 0*.*05, ** p < 0*.*01, *** p < 0*.*001* and *ns* is *non-significant* using Mann–Whitney U tests (Note: mixed linear model with a random intercept for animal ID yielded the same results). In **(E)** the same *p*-value indicators are used with *Wilcoxon signed-rank* tests.

However, this increased vigor in FixGrower-trained animals was accompanied by higher violation rates during fixation growth, indicative of poor task learning (**L:** 21.8 ± 11.6 %, *n*_Session_ = 110,vs. **FG:** 46.1 ± 16.4 %, *n*_Session_ = 110; Mann–Whitney U =1977.5, *p* = 4.78 × 10^−37^; **Fig. 7d**). Crucially, when we introduced the 2-second violation penalty in a single session prior to the probe stages, their violation rate immediately decreased from 47.4 ± 16.5 % to 16.2 ± 5.3% (*Friedman* χ^2^(2) = 13.5, *p* = 0.0011; Day −1 vs. Day 0, *Wilcoxon signed-rank, W*= 0.039, *p*_holm_ = 0.012) and remained consistently low during the subsequent probe session (18.4 ± 8.9%; Day 0 vs. Day 1, *Wilcoxon signed-rank, W*= 0.039, *p*_holm_ = 0.012; **Fig. 7e**). These findings suggest that the elevated violation rates during fixation growth in animals trained with FixGrower were not due to impaired task comprehension, but rather reflected strategic behavior shaped by the minimal consequences of failure.

Correspondingly, the introduction of the penalty significantly increased the cost of failure, markedly slowing post-violation trial initiation times for FixGrower animals, bringing their behavior closer to that of Legacy animals (**L:** median 5.70s, mean ± SD 23.0 ± 58.7s, *n*_trials_ = 2,960 vs. **FG:** median4.47s, mean ± SD 23.5 ± 66.9s, *n*_trials_ = 3,095; *U =* 5.04 × 10^6^, *p* = 7.82 × 10^−12^; **Fig. 7f**). Moreover, we observed no significant difference between cohorts in the number of trials attempted during the probe stages (**L:** 345 ± 77 vs. **FG:** 359 ± 97; Welch’s *t* = 0.544, *p* = 0.595; **Fig. S1a**). Collectively, these results demonstrate that large negative reinforcement early in learning is associated with reduced behavioral vigor in our task, which—when coupled with the trial-dependent structure of the Legacy growth algorithm—likely contributed to slower fixation growth in the Legacy cohort.

We next investigated how metrics of engagement and motivation evolved within individual sessions, given that task difficulty progressed differently for each cohort. In the Legacy curriculum, task difficulty incrementally increases throughout each session, as successful trials progressively raised the fixation requirement. In contrast, the FixGrower curriculum has peak task difficulty at session onset, where animals need to immediately adapt to the new fixation requirement set by the prior session. We thus predicted that engagement would decrease over sessions in Legacy-trained animals, while remaining stable or possibly improving in FixGrower-trained animals.

For Legacy-trained animals to optimally grow fixation duration, they ideally maximize their trial rate and minimize violation rates. We therefore analyzed these two metrics specifically during the first 50 and last 50 trials within sessions containing at least 200 trials as proxies of engagement. Consistent with our predictions, during the fixation growth stages Legacy animals exhibited modest, yet significant increases in violation rates from early to late session (Early: 22.5 ± 11.7 % vs. Late: 25.6 ± 14.6 %, *n*_Session_ = 216; *Wilcoxon signed-rank W* = 7803, *p*_holm_ = 3.22 × 10^−3^; **Fig. 8a**). In contrast, FixGrower animals showed significantly decreased violation rates from early to late session (Early: 66.1 ± 14.1%vs. Late: 35.6 ± 13.7%, *n*_Session_ = 102; *W* = 8, *p*_holm_ = 6.67 × 10^−18^; **Fig. 8a**).

**Figure 8.**
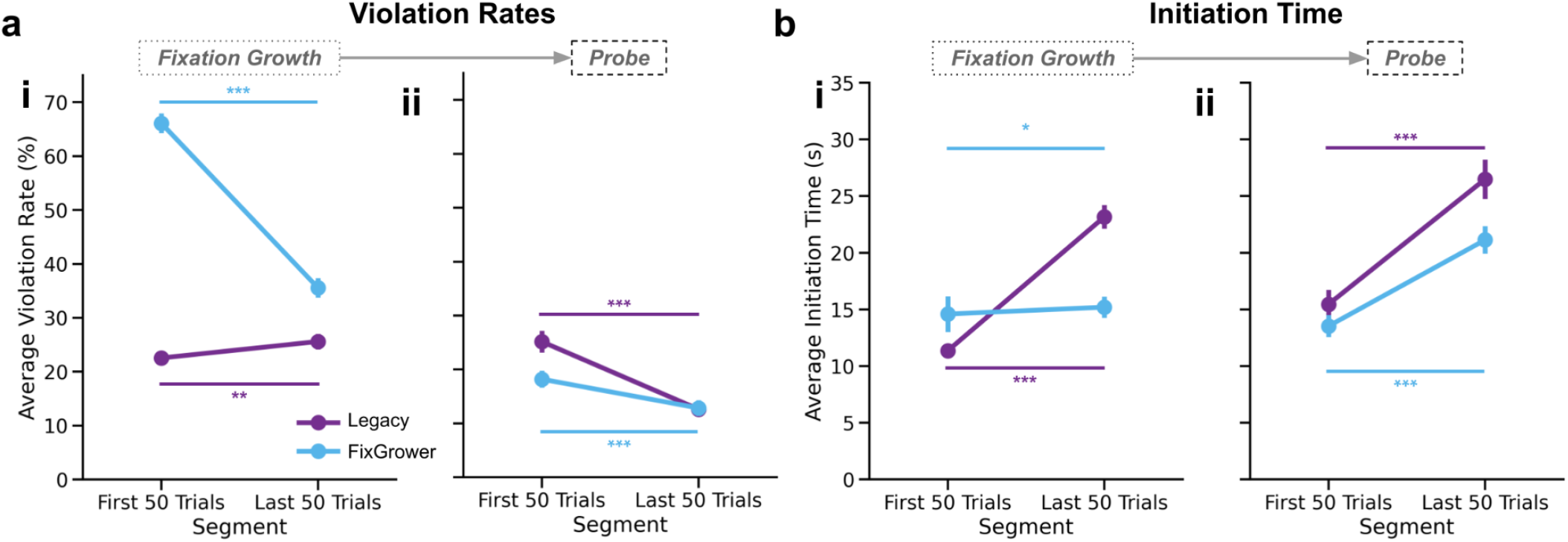
FixGrower-trained animals have more stable within-session engagement than Legacy-trained animals. **(A)(i)** Violation rates comparing the first 50 trials to the last 50 trials within each session during the fixation growth curriculum for each cohort (Legacy: dark purple, FixGrower: light blue). **(A)(ii)** As in **(A)(i)** but during the probe stages when task difficulty was stable and matched across cohorts. **(B)(i)** Trial initiation time following all trial types comparing the first 50 trials to the last 50 trials within each session during the fixation growth curriculum for each cohort. **(B)(ii)** Trial initiation times as in **(B)(i)** but during the probe stages when task difficulty was stable and matched across cohorts. *Statistics:* Circles represent group means and error bars indicate standard error of the mean. Statistical comparisons were conducted within groups (e.g., Legacy early vs. Legacy late) using *Wilcoxon signed-rank* tests. * *p < 0*.*05, ** p < 0*.*01, *** p < 0*.*001* and *ns* is *non-significant* using Holm correction for multiple comparisons.

Similarly, Legacy-trained animals displayed significantly increased trial initiation times (i.e., the time subjects took to start a new trial after the ITI had passed) from early to late session (Early: 11.4 ± 4.08s vs. Late: 23.2 ±12.2s, *n*_Session_ = 216; *W*= 920, *p*_holm_ = 3.96 × 10^−18^; **Fig. 8b**). Whereas, initiation times in FixGrower-trained animals only increased by half a second on average (Early: 14.5 ± 13.9s vs. Late: 15.2 ± 7.39s, *n*_Session_ =102; *W*= 1988, *p*_holm_ = 0.033; **Fig. 8b**).

Critically, these divergent within-session patterns were not observed during the subsequent probe stages, where task difficulty remained stable and matched between cohorts throughout each session. Under these matched conditions, both cohorts demonstrated significantly decreasing violation rates from early to late session (**L Early:** 25.2 ± 12.7%vs. **L Late:** 12.6 ±7.84, *n*_Session_ = 67; *W* =104.5, *p*_*holm*_ *=* 4.91 × 10^−10^; **FG Early:** 18.2 ± 9.53% vs. **FG Late:** 12.8 ± 8.74%, *n*_Session_ = 71; *W* = 508.0, *p*_holm_ = 2.82 × 10^−5^; **Fig. 9a(ii)**), consistent with improving proficiency, and significantly increasing trial initiation times (**L Early:** 15.5 ± 8.79s vs. **L Late:** 26.5 ± 12.5s, *n*_Session_ = 67; *W* = 190, *p*_holm_ = 6.13 × 10^−9^; **FG Early:** 13.5 ± 6.42s vs. **FG Late:** 21.1 ± 8.42s, *n*_Session_ = 71; *W* = 338, *p*_holm_ = 7.20 × 10^−8^; **Fig. 8b(ii)**), consistent with accumulating fatigue or satiety. Taken together, these data indicate that incremental within-session increases in task difficulty are associated with reductions in performance and engagement, factors which likely contributed to the significantly slower fixation growth observed in Legacy-trained animals.

**Figure 9.**
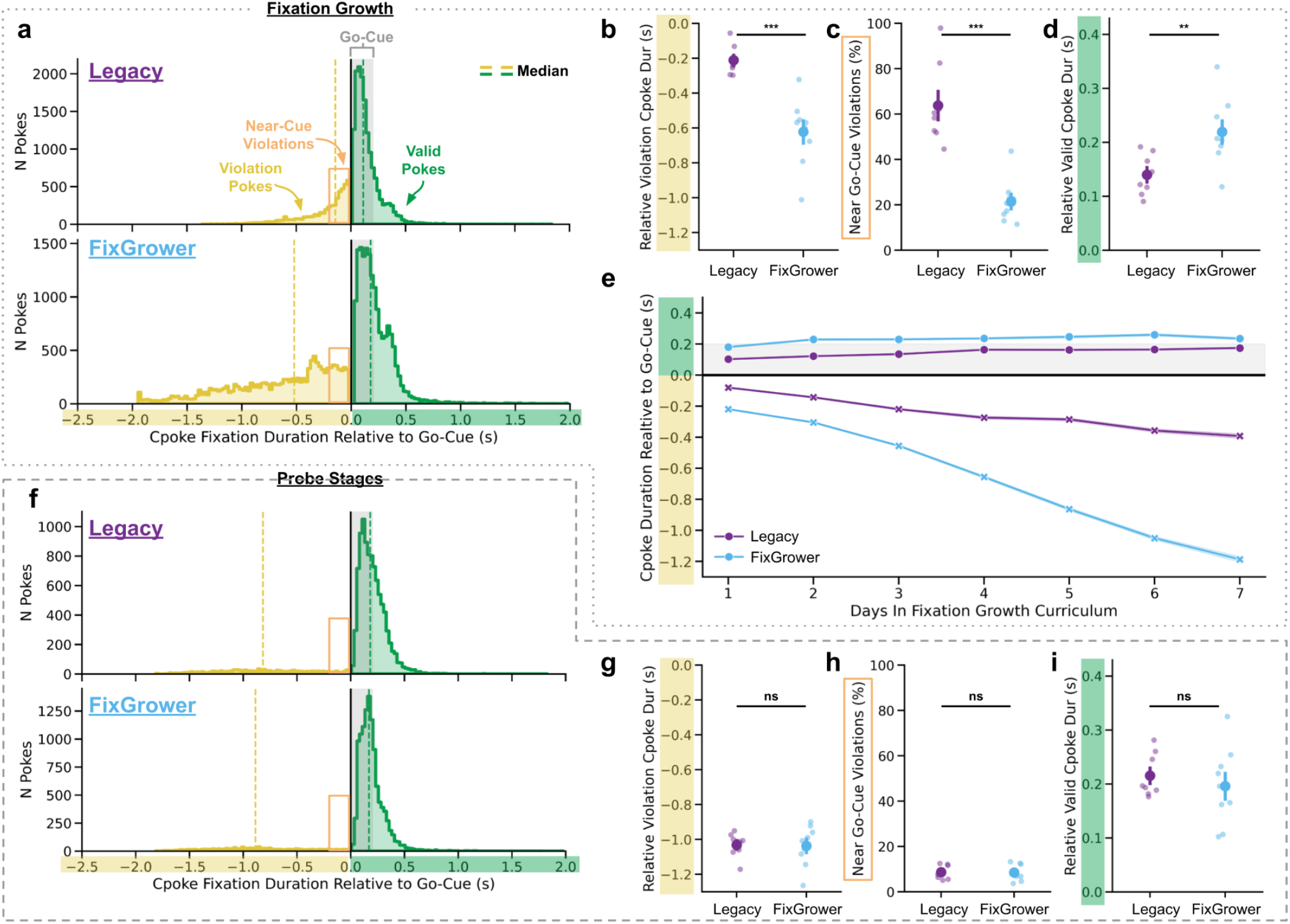
Legacy-trained animals exhibit more impulsive center-poking behavior than FixGrower-trained animals during fixation training. **(A)** Distributions of center-poke durations relative to the go-cue during the first 7 days of the fixation growth curriculum for Legacy (top, dark purple) and FixGrower (bottom, light blue) animals. Histograms show counts of all pokes with valid pokes colored green and violation pokes colored yellow. Dashed lines indicate respective medians. The orange box marks the “near-cue” period, defined as the 200 ms window immediately preceding the go-cue. (**B**) Mean violation poke durations relative to the go-cue for each cohort. **(C)** Percentage of violations occurring within the near-cue period defined in (A). (D) Mean valid poke durations (i.e., post-go-cue) relative to the go-cue for each cohort. (E) Daily averages of valid (green) and violation (yellow) poke times relative to the go-cue across the 7 days of fixation growth. Lines represent cohort means ± SEM. (F) Same as (A), but for the probe stages when curriculum differences were eliminated. (G–I) Same as (B–D), but calculated from probe session data. Statistics: All scatter/point plots display mean ± SEM, with individual animal data points overlaid. * *p < 0*.*05, ** p < 0*.*01, *** p < 0*.*001* and *ns* is *non-significant* using Welch’s t-tests.

Lastly, we examined how differing starting fixation requirements and growth algorithms influenced center-poke timing, with a focus on impulsivity and motor control. The Legacy curriculum began each session with a minimal fixation requirement (10 ms) that incrementally increased after successful trials. While this structure eases animals into fixation, it rewards short, rapid center pokes early on. Moreover, because fixation demands grew only after successful trials, previously rewarded movement patterns could become violations on subsequent trials, potentially disrupting learning through inconsistent reinforcement. In contrast, the FixGrower curriculum employed a stable, longer initial fixation requirement (400 ms) within each session, encouraging sustained fixation from the outset.

Given these structural differences, we hypothesized that Legacy-trained animals would display more impulsive center-poking behavior. We predicted this would manifest as temporal clustering around the moments before the go-cue, which would minimize the temporal difference between rewarded and punished center-pokes, thereby making discrimination of optimal behavior more difficult. To test this, we compared the timing of valid and violation pokes relative to the go-cue during the first week of fixation growth. This period was selected because it matched the time FixGrower-trained animals took to complete fixation growth, allowing for a temporally aligned behavioral comparison. Consistent with our hypothesis, poke-time distributions differed markedly between groups (**Fig. 9a**). Legacy-trained animals violated significantly closer to the go-cue compared to FixGrower-trained animals (**L:** mean ± SD −0.211 ± 0.008s, median : −0.225s, *N* = 8; **FG:** −0.622 ± 0.193s, median : −0.569s, *N* = 9; Welch’s *t* = 5.81, *p* = 1.12 ×10^−4^; **Fig. 9b**). Additionally, a significantly higher fraction of violations in Legacy-trained animals occurred within the immediate 200 ms preceding the go-cue (**L:** 63.7 ± 17.7%, *N* = 8; **FG:** 21.4 ± 9.58%, *N* = 9; Welch’s *t* = 6.03,p = 1.04 × 10^−4^; **Fig. 9c**), suggesting these animals were anticipating rather than reacting to the go-cue—likely a result of the curriculum differences highlighted above that fostered premature responses.

For valid pokes, FixGrower-trained animals exhibited significantly longer durations post-go-cue compared to Legacy-trained animals (**L:** 0.140 ± 0.038s, median : 0.133s, *N* = 8; **FG:** 0.219 ± 0.062s, median : 0.207s, *N* =9; Welch’s *t* =−3.23, *p* = 6.27 × 10^−3^; **Fig. 9d**), and these group differences were evident from the first day of fixation growth (Welch’s *t, W*= 0.039, *p*_bonferroni_ < 0.001 for all days, valid and violation between cohorts; **Fig. 9e**). To assess whether these differences could be attributed to the initial 20-trial “warm-up” in the Legacy curriculum, during which fixation requirements are closest to the go-cue, we reanalyzed the data excluding these trials. Results remained consistent, confirming that observed differences were not significantly influenced by early-session structure (**Fig. S5a-d**). Repeating analyses across the entire fixation growth phase (not just the first week) showed that Legacy-trained animals’ valid poke durations gradually increased, eventually approaching those of FixGrower-trained animals (**L:** 0.198 ± 0.047s, median : 0.192s, *N* = 8; **FG:** 0.216 ± 0.0068s, median : 0.216s, *N* = 9; Welch’s *t =* −0.650, *p* =0.526), whereas the violation poke metrics remained unchanged (**Fig. S5e-i**).

During the probe stages, all group differences in poke-timing disappeared (**Fig. 9f-i**). Legacy-trained animals exhibited poke distributions, average violation timing (**L:** −10.31 ± 0.070s, median : −1.010s, *N* = 8; **FG:**−1.036 ± 0.116s, median : −1.008s, *N* = 9; Welch’s *t* = 0.114, *p* = 9.10), near-go-cue violations (**L:** 8.68 ± 3.04%, *N* = 8: **FG:** 8.48 ± 3.49%, *N* = 9; Welch’s *t* =0.131, *p* =0.897) and valid poke durations (**L:** 2.15 ± 0.041s, median : 0.195s, *N =* 8; **FG:** 0.196 ± 0.071s, median : 0.196s, *N* = 9;Welch’s *t* = 0.705, *p* = 0.493) similar to those of FixGrower-trained animals. Notably, FixGrower-trained animals showed consistent timing behavior across both the fixation growth and probe stages, indicating a stable motor strategy throughout training.

These findings suggest the Legacy curriculum placed animals at an initial disadvantage: by rewarding brief pokes and introducing shifting contingencies, it likely fostered impulsive habits and delayed the formation of stable fixation behavior. In contrast, the FixGrower curriculum provided consistent reinforcement for sustained fixation throughout training. This led to more deliberate poking, which directly contributed to increased training speed, as longer valid pokes led to larger growth increments for these animals.

In summary, the three curriculum features—*penalty timing, growth algorithm*, and *starting fixation requirement*—exerted multiple, compounding effects on behavior. Together, these factors compounded to hinder learning in the Legacy curriculum while synergistically enhancing training efficiency and behavioral stability in FixGrower curriculum by modulating vigor, engagement and motor control.

## Discussion

Perceptual decision-making tasks require animals to learn both stimulus–response rules and structured motor behaviors. One such behavior is center-port fixation, where a freely moving animal must initiate and maintain a nose poke at a central port for a specified duration. Although fixation is a common task requirement, the procedures used to train this behavior vary widely and often require substantial time, presenting an opportunity for optimization and standardization. In response to this, we developed a novel curriculum for shaping fixation behavior in rodents, designed to minimize both training duration and premature-movement fixation errors termed “violations”. Our approach, called “FixGrower”, differed from commonly used methods in three key ways: it began with a long initial fixation duration of 400 ms, it increased the requirement adaptively across days rather than trials, and it delayed the introduction of violation penalties until the animal reached the target fixation duration.

To evaluate the effectiveness of this curriculum, we trained two cohorts of rats to reach a 2-second fixation target. One cohort (*N = 8*) followed a curriculum commonly used in our lab and similar to those in the field which we called (“Legacy”, while the other (*N = 9*) was trained using the new FixGrower approach (**Fig. 1**). FixGrower-trained animals reached the target duration significantly faster than Legacy-trained animals, representing a 61% reduction in training time on average (**Fig. 2c**).

We then assessed violation rates under both stable and variable fixation conditions and found that FixGrower-trained animals maintained low violation rates, averaging 13.2% (**Fig. 3a**). Moreover, violation rates and training times were positively correlated in the FixGrower curriculum, providing experimenters with a straightforward metric for predicting cohort performance and setting training cutoffs (**Fig. 3c**). These improvements proved to be robust: in a subset of animals trained for over 100 days following FixGrower completion, violation rates remained consistently low, averaging 9.9% (**Fig. 4**).

To confirm that these low violation rates were not the result of a simplified task structure, we trained a new cohort of rats on a sound discrimination task following FixGrower training (**Fig. 5a-b**). Even with the added requirement of processing task-relevant stimuli during fixation, violation rates remained low at an average of 11.2% (**Fig. 5e**), which is notable given recent findings that lower violation rates are associated with improved stimulus rule performance (Reuschenbach et al., 2023).

To our knowledge, only one study has reported faster fixation acquisition than our approach. Odoemene et al. 2018 trained mice to reach a 1.1-second fixation duration within 1–2 sessions by delivering a small water reward (0.5 μl) at the center port during shaping. However, without this mid-trial reward, their procedure, which was structurally similar to our Legacy curriculum, required 10–12 sessions to reach the same target. While their strategy is highly effective, it introduces both logistical and interpretive challenges: introducing center-port water solely for fixation growth adds complexity to behavioral setups, and rewarding animals prior to stimulus presentation may alter motivational state, affect neural activity, and/or complicate downstream learning.

To evaluate whether our approach could provide a practical alternative, we applied the FixGrower curriculum to a cohort of mice and trained them to a 1.1-second fixation duration. These animals reached the target in just 3–5 days (**Fig. 6b**), and a subset of these animals progressed to fixation durations beyond 2 seconds within 9 days (**Fig. S4a**). Importantly, their violation rates were equivalent to or lower than those published in the field (Ashwood et al., 2022; Cruz et al., 2022; Jaramillo & Zador, 2014; Odoemene et al., 2018) (**Fig. 6d**). These results underscore the utility of the FixGrower curriculum as a robust, scalable solution for shaping fixation behavior in both mice and rats—achieving high performance while preserving standard equipment and task structure.

### Rationale

Understanding why the FixGrower curriculum accelerated fixation training is essential for refining future protocols and contextualizing these results within broader frameworks of learning. To this end, we compared behavioral dynamics during the differing fixation growth stages with those in the probe stages, where task parameters were held constant across experimental cohorts (**Figs. 7-9**). All three curriculum elements—violation penalty introduction, fixation growth algorithm, and initial fixation requirement—appeared to shape learning by modulating motivation, engagement, and motor control. Moreover, we draw on theories from operant conditioning, reinforcement learning, and curriculum learning to interpret these findings.

During fixation growth, Legacy-trained animals completed 30% fewer trials per session than FixGrower-trained animals, despite achieving similar numbers of rewarded trials (**Fig. 7a-b**). This discrepancy was driven by slower trial re-initiation after violations in Legacy-trained animals (**Fig. 7c**); a pattern we hypothesized was influenced by the higher cost of failure imposed by the timeout penalty present only in the Legacy curriculum. When the penalty was introduced in the FixGrower curriculum after reaching target fixation, violation rates dropped markedly and trial initiation times slowed to Legacy-like levels (**Fig. 7e-f**). This behavioral inflection supports theories from operant conditioning suggesting that excessive early punishment can reduce motivation (Skinner, 1965), and is consistent with findings that animals re-engage more quickly post-failure when expected reward is high (Mah et al., 2023). Taken together, these results show that early shaping should prioritize rapid re-attempts post-failure and preserve sizable penalties until later in training so as to keep motivational drive high and facilitate initial learning.

Notably, decreasing vigor and motivation among Legacy-trained animals emerged not only across sessions but within sessions as well. Over the course of individual sessions, these animals exhibited increasing violation rates and slower trial initiation, consistent with reduced engagement (**Fig. 8a(i), b(i)**). We hypothesize this arose from the Legacy curriculum’s incremental fixation growth algorithm, which steadily increased task difficulty each trial, likely contributing to cumulative fatigue. Since both trial count and violation rate determine the growth magnitude in the Legacy curriculum, this within-session disengagement leads to slower fixation growth. In contrast, FixGrower-trained animals maintained stable engagement over time (**Fig. 8a(i), b(i)**), and both groups showed comparable behavior during probe sessions when difficulty was static (**Fig. 8a(ii), b(ii)**), reinforcing the idea that increasing task demands negatively impacted the Legacy cohort.

We further hypothesized that the variable demands of the Legacy curriculum impaired motor control. Because fixation duration increased only after successful trials, a fixation hold that was previously rewarded could be penalized on the very next attempt. This inconsistency mirrors the challenge of learning under non-stationary targets in reinforcement learning, where shifting objectives can destabilize learning by introducing ambiguity into the action–outcome mapping (Sutton & Barto, 2018). From this perspective, each increase in fixation requirement after a valid trial effectively changed the rule the animal was attempting to learn. Behaviorally, this was reflected in Legacy-trained animals exhibiting a smaller separation in timing between failed and successful pokes (**Fig. 9a-e**), and a greater concentration of violations immediately preceding the go-cue—suggesting anticipatory, impulsive behavior rather than deliberate, rule-based control.

One well-established solution to the problem of non-stationary targets in artificial learning systems is the use of a separate, slowly-updated target network (Mnih et al., 2015). This strategy stabilizes learning by keeping the goal (“target”) fixed for a period of time, allowing the agent to learn from consistent feedback instead of constantly adjusting to shifting objectives. Analogously, the FixGrower curriculum achieved behavioral stability by holding the fixation requirement constant within each session—effectively implementing a fixed “target” rule over a session. This provided animals with a consistent behavioral contingency to learn from, thereby promoting clearer reinforcement signals. Just as deep Q-learning benefits from steady targets, keeping task difficulty constant within sessions appears important for sustaining engagement and supporting efficient learning.

Finally, we propose that beginning with an extremely short (10 ms) fixation duration in the Legacy curriculum may have reinforced exploitation, non–goal-directed behaviors that slowed learning. The underlying assumption of the Legacy design—that fixation is a continuum from short to long holds—may not align with how animals conceptualize these actions. Instead, we hypothesize that short pokes and sustained holds constitute qualitatively distinct behaviors. By initially reinforcing rapid pokes, the Legacy curriculum may have inadvertently encouraged habits that later conflicted with the task’s demands. This is reflected in the Legacy-trained animals’ impulsive poke profiles and shorter valid hold durations following the go-cue (**Fig. 9a-e**). As these animals progressed to longer fixation requirements—and later into the probe stages—their post–go-cue hold durations increased (**Fig. S5e-i**), suggesting that they gradually abandoned the initial strategy of rapid, impulsive poking in favor of more deliberate fixation holds.

By contrast, FixGrower’s initial fixation requirement was critically set to exceed the duration of a quick nose flick (∼250 ms), yet remain attainable. This structure increased the salience of the goal behavior by only rewarding meaningful approximations of the desired behavior. This approach aligns with the curriculum learning principle of using carefully chosen examples that are informative yet achievable (Bengio et al., 2009). By shaping behavior through reinforcement of stable, goal-relevant responses, the FixGrower design facilitated more deliberate motor control and efficient learning.

Together, these findings highlight how subtle structural choices in early training can shape the trajectory of learning by influencing the clarity, consistency, and magnitude of feedback. The Legacy curriculum imposed high penalties early and offered less stable reinforcement contingencies, leading to reduced engagement and noisier learning. In contrast, the FixGrower curriculum supported behavioral shaping through milder early consequences and clearer, more consistent reinforcement.

### Limitations and future directions

A key limitation of this study is the inability to isolate the specific contributions and interactions of the three curriculum features that contributed to improved learning efficiency using the FixGrower curriculum. Future work could address this by conducting a systematic “knockout” analysis, in which animals or theoretical models are trained with subsets of the features to examine how behavioral metrics are affected.

A second limitation concerns task generalization across different delay durations. Some paradigms, such as those used in Akrami et al. 2018, incorporate delays exceeding 2 seconds, which may not be efficiently supported by overnight growth alone. To accommodate longer durations, we recommend supplementing overnight growth with intra-session increases. For example, by incrementing fixation duration every 100–150 trials once 1–2 seconds are reliably achieved. Experimenters should monitor motivational levels closely and consider implementing 5–10 “warm-up” trials to support adaptation to extended delays.

Third, not all behavioral tasks require or benefit from an auditory go-cue (for example: Brunton et al., 2013; Kopec et al., 2024). While we view the auditory cue as an important feature for supporting this fixation task, further investigation is needed to identify alternative salient cues or to develop strategies for phasing out the cue once fixation behavior is stable.

In summary, we present FixGrower: a novel curriculum-based approach for training center-portfixation in rodents that significantly reduces training duration while maintaining low rates of premature movement. Our results are robust across tasks and species, and grounded in principles from operant conditioning, curriculum learning, and reinforcement learning. We anticipate that this work will be of broad utility for the efficient design and implementation of behavioral training protocols in psychology and neuroscience research.

## Methods

### Animal Subjects

#### Rats

We used male Long-Evans rats (*Rattus norvegicus*) between the ages of 4 and 7 months for this study. All rats were pair housed and had *ad libitum* access to standard rodent chow. Rats did not begin water restriction and training until their weight exceeded 250 g. For at least one week prior to training, animals were brought to the training room daily for weighing. Once on restriction, water was available during the behavioral session (28–60 μL for each correct trial depending on animal and stage) and for 1 hour at the end of the session. Rats were removed from water restriction if they showed signs of illness, failed to consume at least 3% of their body weight in water over three consecutive days, lost greater than or equal to 10% body weight in 24 hours, lost greater than or equal to 20% body weight over 2 weeks or less, or exhibited a statistically significant trend of greater than or equal to 1.5% weight loss per week over 30 days.

#### Mice

We used male and female *B6*.*CAST-Cdh23Ahl+Kjn* mice (The Jackson Laboratory, stock #002756). Animals were group-housed by sex in standard ventilated cages (2–5 animals per cage) under a 12-hour light/dark cycle with *ad libitum* access to standard rodent chow.

Upon arrival, animals were acclimated to the housing environment and handled using a non-aversive tunnel-based approach to reduce stress, as described by Gouveia & Hurst, 2013. Briefly, a tunnel (Kent Scientific) was placed in each home cage to allow voluntary entry, and over the course of a week, mice were gently habituated to being picked up first via the tunnel and then by hand.

Once mice were at least 8 weeks old, seven days prior to the start of behavioral training, they were weighed to establish a baseline body mass. At this time, they were transitioned to 2% citric acid (Thermo Scientific Chemicals, CAS: 77-92-9) to begin water restriction (Urai et al., 2021). From this point forward, mice were weighed daily, and a daily Body Condition Score (BCS) assessment was conducted to monitor animal welfare (Burkholder et al., 2012). Following the one-week acclimation to water restriction, animals began behavioral training. During behavioral sessions, water was available as a reward for correct trials (3–10 μL per trial, depending on the animal and training stage). If a mouse failed to consume at least *max(*1 mL, 4% of baseline body weight) it was supplemented with the remaining water to reach this threshold post training session. On non-training days, animals were provided *ad libitum* access to 2% citric acid water. If an animal did not train for seven consecutive days, it received 1 hour of *ad libitum* access to water. If an animal’s weight dropped more than 10% below baseline, *ad libitum* water access was reinstated until the weight stabilized. Additionally, if an animal lost more than 1 g of body mass within 24 hours, dehydration was assessed via skin turgor. If dehydration was indicated, the animal was supplemented with free water.

### Behavioral Training

#### Apparatus

Behavioral training took place in custom-designed operant conditioning boxes housed within sound-attenuating chambers equipped with active ventilation. Each training box featured three conical nose ports arranged in a horizontal row (Left – Center – Right; **Fig. 1a**). Planar tweeters (Bohlender Graebener Neo3W) were mounted above the left and right nose ports to deliver auditory stimuli. Each port was equipped with a controllable white LED for visual cues and an infrared beam to detect nose pokes.

The left and right ports were fitted with solenoid valves (Lee Valve Company, LHDA1231115H) connected to sipper tubes that dispensed distilled water as a reward, sourced from a 10-gallon gravity-fed tank. The center port had an enlarged opening to facilitate nose fixation, allowing rats to maintain position comfortably.

All hardware was controlled by a Bpod 2.3 system (Sanworks) interfaced with a Windows computer running Bcontrol, a custom MATLAB-based behavioral training software (https://tinyurl.com/BrodyBcontrol, https://tinyurl.com/BrodyExperPort). Auditory stimuli were generated via an Analog Output Module (Sanworks), amplified by a mini class D stereo amplifier (Leapai LP-2020AD), and delivered through the planar tweeters.

Mouse training boxes were similarly configured but downsized in proportion to the animals’ size. In the mouse setup, water reward was supplied from a 100 mL syringe located within the chamber and refilled after each session.

### Training Overview

Eighteen rats participated in the primary fixation training study, divided into two cohorts that experienced differing fixation growth curricula: Legacy (*N = 9*) and FixGrower (*N = 9*). Rats were pair-housed such that each pair included one Legacy and one FixGrower animal. Water restriction and training commenced once both animals in a pair exceeded 250 g body weight.

Training sessions were conducted daily (7 days per week) and lasted between 1.5 and 3 hours per session. With few exceptions (e.g., equipment failure), each animal was trained in a consistent behavioral box throughout the experiment. An effort was made to distribute training such that both cohorts were represented across the available boxes over the course of the experiment. For example, if in one session a Legacy rat trained in Box #10 and a FixGrower rat in Box #20, a subsequent session might place a Legacy rat in Box #20 and a FixGrower rat in Box #10. This strategy was implemented to help mitigate any potential box-specific effects. All animal handling—including weighing, training, and health checks—was performed by technicians blinded to the experimental conditions. The only experimental variable differing between cohorts was the specific structure of the fixation growth training curricula.

An additional cohort of 13 rats trained on a sound discrimination task followed the same training structure and procedures. The mouse cohort (*N = 5*) also adhered to this protocol with two exceptions: sessions lasted 1 hour, and training began once mice were at least 8 weeks old and had been on 2% citric acid for at least seven days.

### Curricula

#### Overview of Stages for Fixation Experiment

Automated behavioral training was conducted using Bcontrol, a custom MATLAB-based software package (MathWorks, R2019b). A unified protocol file (FixationGrower) controlled the overall task structure and parameters for all animals. Stage-specific training progressions were governed by customized Training Section files, which defined the curricula and were the only components to differ between the experimental cohorts. Animals progressed through these three main experimental phases (**Fig. 1b, Table S1**):

1. A **side poke curriculum** to facilitate reward learning and habituation.
2. A **fixation growth curriculum**, either “Legacy” or “FixGrower.”
3. A set of **“probe” evaluation** stages following completion of fixation growth.

Each training stage is described in detail in the subsequent methods sections. Importantly, the only difference between curricula lies within the fixation growth stages (Stages 5–8 in **Table S1**, highlighted in red); all other training phases and parameters were identical across cohorts. For a detailed account of the underlying state-machine logic implemented in the protocol, refer to **Fig. S3.6** and **Fig. S3.7**.

#### Shared: Side Poking

The first 4 training stages are shared between cohorts and serve the purpose of habituating the animals to the behavior box and water reward while ensuring they are motivated to perform trials (**Table S2**).

##### Stage 1– Learn Left (L) Poking

The structure for Stage 1 trials is the following: go-cue (200 ms white noise at 70 dB, bilateral) → L side-port LED on → L side-port water seeded (30%, 18 μL (mouse: 3 μL)) → wait for animal to poke in L port (max 8 s) → animal pokes in L port → L side-port remaining water delivered (70%, 42 μL (mouse: 10 μL)) & LED turned off → trial inter-trial interval (ITI) sampled from a normal distribution with mean 40 s, standard deviation 5 s. If the animal does not poke in the left port during the wait-for-poke period it is considered a “no answer” trial. In this case, the remaining reward is not delivered, the L side light turns off, and the trial moves into the ITI period. On the trial following a no answer trial, water is not seeded in the L port (we assume it still remains from the previous trial). If the animal pokes in the incorrect port (right) during the wait-for-poke period, there is no penalty and they can immediately retry within the same wait period. To complete this stage, animals must complete 40 trials and have a no answer rate < 15%. They can move to the next stage within the same session. If the animal does not complete this stage on the current day’s session, and they have yet to complete Stage 2, they will move into Stage 2 for the following session such that they alternate between Stage 1 and Stage 2 over days. If they have completed Stage 2, they will stay in Stage 1 until completed before moving to Stage 3. →

##### Stage 2– Learn Right (R) Poking

The structure for Stage 2 is identical to that for Stage 1 except that the right (R) port is used. Additionally, the same stage completion logic follows such that if they fail to complete Stage 2 within the session, they will return to Stage 1 on the following day if Stage 1 has yet to be completed. If Stage 1 is completed, they will remain in Stage 2 until completed. Once Stage 1 and Stage 2 are *both* completed, the animal will move into Stage 3.

##### Stage 3– Learn Left & Right Poking in Blocks

In Stage 3, the animals will alternate between 15-trial blocks of left and right poking. The structure for Stage 3 trials is the following (assume in L block): go-cue sound → L side-port LED on → wait for animal to poke in L port (max 8 s) → animal pokes in L port → L side-port water reward delivered (100%, 45 μL (mouse: 6 μL)) → trial ITI sampled from a normal distribution with mean of 25 seconds, standard deviation of 3 seconds. In this stage, the ports are no longer seeded with water. If the animal pokes incorrectly during the wait-for-poke period, this is considered an “error” trial. Similar to a no answer trial, the side LED turns off immediately and the trial moves into the ITI period. To complete this stage, animals must do at least 90 trials (6 blocks) with > 87% accuracy and a no answer rate < 20%. Note that accuracy is only computed on trials that were answered and stage advancement only occurs at the end of the session (i.e., overnight) for this stage and all future stages.

##### Stage 4– Learn Left & Right Poking in Blocks to Random

In Stage 4, the animals will alternate between 15-trial blocks of left and right poking until they reach 70 trials. Then, each trial will be randomly drawn with an anti-biasing procedure which dynamically adjusts trial probabilities to counteract subject side-biases by considering past trial outcomes. It uses an exponential kernel with a decay parameter, *τ* (default = 30 trials), to estimate recent performance. It then modifies the probability of selecting left or right trials based on this history. The adjustment strength is controlled by *β* (default = 3), which is initialized with a linear warm-up over *τ* trials. The value of *β* is adjusted on a per-animal basis and ranged from 3 to 4.5 in our experiments.

The only adjustments to the trial structure from Stage 3 are the following: the water reward is 30 μL (mouse: 3 μL), and the ITI is sampled from a normal distribution with mean of 15 seconds, standard deviation of 3 seconds (**Table S2**). To complete this stage, animals must do at least 175 trials in a session with > 80% accuracy and < 20% no answer rate. Animals will then move into Legacy or FixGrower curriculum depending on the experiment.

#### Varied: Fixation Growth

The core of this experiment is a comparison between two training curricula designed to shape motor fixation behavior. Animals were trained to reach a target fixation duration of 2 seconds using one of two curricula: Legacy or FixGrower. The curricula differ in three key features: the initial fixation duration, the algorithm governing fixation growth, and the penalty for fixation violations (**Table S3**; **Fig. 1c-e**).

Aside from these differences, both curricula follow the same overall stage progression, as outlined in **Table S4**. In the additional rat and mouse experiments presented in Figures 5 and 6, the FixGrower curriculum was used, with slight modifications to stage 8 that are described in the corresponding sections.

#### Legacy Fixation Growth (Stages 5–7)

Animals in the Legacy experimental cohort progressed through three structured stages to learn the center poke fixation-to-side poke reward motor sequence. These stages were adapted from standard shaping procedures used both in our lab and more broadly in the field (Akrami et al., 2018; Constantinople et al., 2019; Mah et al., 2023; Schmid et al., 2024; Scott et al., 2015). Specifically, they closely follow the “Grow Nose Poke” curriculum described in Akrami et al. 2018, with the primary modification being that the daily warm-up stage was performance-based rather than fixed. This adjustment was made to reduce early-stage dropout, which was frequently observed anecdotally due to animals disengaging from the task.

##### Legacy Stage 5– Center to Side, Blocks with Fixation Growth

In Legacy Stage 5, the animals are required to first poke in the center-port for a required fixation period before receiving water reward localized by LED light in one of the side-ports, which alternates in blocks of 15 trials. The structure for these trials is the following (assume in L block): Center-port LED indicates trial availability → animal pokes & holds nose in center → go-cue sound plays & Center LED off → L LED on & wait-for-poke period (max 8 s) → animal pokes in L port → L side-port water reward delivered (30 μL, mouse: 3 μL) → trial ITI sampled from a normal distribution with mean of 5 seconds and standard deviation of 2 seconds. If the animal leaves the center-port before the go-cue plays, or if the animal concurrently pokes in a side-port during fixation, the trial is considered a “violation.” In this case, the center-port LED stays on only for the intended fixation period, the go-cue does not play, and there is a 2-second penalty timeout. Once the timeout is complete, the trial moves into the ITI period. If the animal stays in the center-port until the go-cue (i.e., a valid trial) but then pokes in the unlit side-port, it is an error trial. If the animal does not poke during the wait-for-poke period, it is a no answer trial. For both error and no answer trials, the side LED turns off immediately and the trial moves into the ITI period.

For the first session in this stage, the starting fixation duration is 0.01 s. After a valid trial, the required fixation duration grows by *max(*1 ms, 0.1%). In other words, the fixation grows linearly by 1 millisecond until the fixation duration exceeds 1 second, and then grows exponentially by 0.1% of the current duration. Assuming the animal performed valid trials in the first session, for the second session—and all remaining sessions in this curriculum—there is a 20-trial fixation warm-up starting at 10 milliseconds and growing toward the previous session’s final fixation value. The growth size in this warm-up block is computed as: 

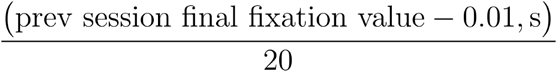

The same min-growth logic applies in the warm-up block: growth only occurs after valid trials. To complete this stage, animals must perform at least 200 trials in a session with > 85% accuracy and < 10% no answer rate. Note that accuracy is computed only on valid trials.

##### Legacy Stage 6– Center to Side, Blocks to Random with Fixation Growth

Legacy Stage 6 is identical to Legacy Stage 5 except that the blocked L/R trials end at trial 70 and then become randomly sampled with the anti-biasing procedure (as described in Stage 3). All other trial structures (fixation, violation rules, reward size, ITI distribution) remain the same. To complete this stage, animals must perform at least 250 trials in a session with > 85% accuracy and < 10% no answer rate.

##### Legacy Stage 7– Center to Side, Random with Fixation Growth

Legacy Stage 7 is identical to Stages 5 and 6 except that all trials are randomly sampled with anti-biasing from the start. To complete this stage, animals must have grown to the final fixation target duration (2 seconds) and performed at least 200 trials in that session, with > 85% accuracy and < 10% no answer rate. Note that only the final trial in the session needs to reach greater than or equal to 2-seconds fixation duration for the stage to be considered completed. From this stage, Legacy animals move directly into probe Stage 9.

#### FixGrower Fixation Growth (Stages 5–8)

Animals in the FixGrower experimental cohort progressed through 4 structured stages to learn the center poke fixation-to-side poke reward motor sequence as described below.

##### FixGrower Stage 5– Center to Side, Blocks with Fixation Growth

Identical to Legacy Stage 5, FixGrower Stage 5 requires animals to initiate a trial by poking into the center-port and maintaining fixation for a specified duration, after which a water reward is delivered at one of the side-ports, indicated by an illuminated LED. The rewarded side alternates in blocks of 15 trials. The structure for these trials is the following (assume in L block): Center-port LED indicates trial availability → animal pokes & holds nose in center → go-cue sound plays & Center LED off → L LED on & wait-for-poke period (max 8 seconds) → animal pokes in L port → L side-port water reward delivered (30 μL, mouse: 3 μL) → trial ITI sampled from a normal distribution with mean of 5 seconds and standard deviation of 2 seconds. If the animal leaves the center-port before the go-cue plays, or the animal concurrently pokes in a side-port during fixation, the trial is considered a violation.

This stage marks the beginning of divergence between the two curricula. In FixGrower Stage 5, if a fixation violation occurs, the center LED remains illuminated indefinitely, indicating that the trial is still active and allowing the animal to immediately retry. The center LED is only extinguished when the animal successfully maintains fixation until the go-cue is played. If the animal holds fixation until the go-cue (i.e., a valid trial) but then pokes into the unlit side-port, the trial is classified as an error. If the animal fails to initiate a trial during the wait-for-poke window, it is labeled a no answer trial. In both error and no answer trials, the side LED turns off immediately, and the trial transitions into the ITI period.

In the first session of this stage, the required fixation duration is set to 0.4 seconds. For all subsequent sessions, the fixation requirement is updated based on the animal’s performance in a given session. If the animal had a bad session—defined as fewer than 20 valid trials—and has not yet begun growing (i.e., the required fixation duration is less than or equal to 0.4 seconds), the fixation requirement for the following session is reduced by 50 milliseconds to lower the entry barrier for initial trials. If the animal has a good session—defined as more than 50 valid trials—and the current fixation requirement is still below the 2-second target, the next day’s requirement increases to match the average center-poke duration on valid trials from the current session, with an upper limit of 2.5 seconds. However, if the animal has a bad session after it has already begun growing (i.e., the required fixation duration exceeds 0.4 seconds), a reset rule is applied: the next day’s fixation requirement is reduced to the midpoint between today’s requirement and the average duration of failed fixation attempts. If the animal’s performance does not qualify as either a “good” or “bad” session—defined as completing between 21 and 50 valid trials—the fixation duration remains unchanged. To complete this stage, animals must complete at least 200 trials in a single session with an accuracy greater than 85% and a no answer rate below 10%.

##### FixGrower Stage 6– Center to Side, Blocks to Random with Fixation Growth

FixGrower Stage 6 is identical to FixGrower Stage 5 except that the blocked L/R trials end at trial 70 and become randomly sampled with anti-biasing. To complete this stage, animals must perform at least 250 trials in a session with greater than 85% accuracy and less than 10% no answer rate.

##### FixGrower Stage 7– Center to Side, Random with Fixation Growth

FixGrower Stage 7 is identical to FixGrower Stages 5 and 6, with the exception that all trials are randomly sampled using anti-bias logic. To complete this stage, animals must perform at least 200 trials in a single session with a fixation duration exceeding the target, while maintaining greater than 85% accuracy and a no answer rate below 10%. For the primary fixation curricula experiment, the target was 2 seconds. For the rat sound discrimination experiment, the target was 1.45 seconds. For the mouse fixation experiment, the target was 1.1 seconds. Unlike Legacy animals, FixGrower animals proceed to one additional stage before entering the probe phase.

##### FixGrower Stage 8– Violation Penalty On

FixGrower Stage 8 maintains the same trial structure as Stage 7, with the addition of a post-violation penalty timeout and a defined inter-trial interval. During the fixation period, a 150-millisecond free-retry window is implemented: if the animal enters and exits the center-port within this window, the trial immediately restarts without being counted as a violation. After this window, if the animal breaks fixation—either by exiting the center-port before the go-cue plays or by poking a side-port during fixation—the trial is classified as a violation. In such cases, the center-port LED remains on only for the originally intended fixation duration, the go-cue is not played, and a 2-second penalty timeout occurs followed by the ITI sampled from a normal distribution with mean of 5 seconds and standard deviation of 2 seconds.

Because animals have already reached the target fixation duration by this stage, the requirement is no longer increased. Instead, the required fixation duration is sampled independently for the session from an exponential distribution with a minimum of 1 second, a maximum of 2 seconds, and a time constant (*τ*) of 1.2 seconds. In the fixation curriculum experiment, after a single session in Stage 8, animals proceed to the fixation evaluation probe stages (Stage 9), regardless of performance.

#### Shared: Fixation Evaluation Probe Stages

There are two probe stages that all fixation experiment animals pass through once growth stages are complete to assess fixation performance, with each stage being 5 sessions in length.

##### Fixation Evaluation Stage 9– Probe Stable Fixation Delay

In Stage 9, trials are randomly sampled L/R with anti-biasing and have a stable fixation duration of 2 seconds. The trial structure is as follows (assuming L trial): Center-port LED indicates trial availability → animal pokes & holds nose in center (2 seconds) → go-cue sound plays & Center LED off → L LED on & wait-for-poke period (max 8 s) → animal pokes in L port → L side-port water reward delivered (30 μL, mouse: 3 μL) → trial ITI sampled from a normal distribution with a mean of 4 seconds and standard deviation 1 second.

During the fixation period, a 150 milliseconds free-retry window is implemented: if the animal enters and exits the center-port within this window, the trial immediately restarts without being counted as a violation. After this window, if the animal breaks fixation—either by exiting the center-port before the go-cue plays or by poking a side-port during fixation—the trial is classified as a violation. In such cases, the center-port LED remains on only for the originally intended fixation duration, the go-cue is omitted, and a 2 seconds penalty timeout follows. This approach allows us to isolate violation trials that reflect active disengagement from the port, rather than brief exploratory movements as the animal settles into position.

If the animal stays in the center-port until the go-cue (i.e., a valid trial) but pokes in the unlit side-port, it is an error trial. If the animal does not poke during the wait-for-poke period, it is a no answer trial. For both of these trial types, the side LED turns off immediately and the trial moves into the ITI period. After 5 days in this stage, the animal moves into Stage 10.

##### Fixation Evaluation Stage 10– Probe Random Fixation Delay

Stage 10 is identical to Stage 9 except that the fixation duration is randomly sampled each trial from an exponential distribution with a minimum of 1 second, a maximum of 2 seconds, and a time constant (*τ*) of 1.2 seconds. After 5 days in this stage, the animal has completed the fixation experiment.

#### FixGrower Animals Post-Probe

A subset (*N = 8*) of FixGrower animals continued training after the end of the experiment in Stage 11 (**Fig. 4a**). This stage was identical to Probe Stage 10 except that the fixation duration was set at the beginning of each session by sampling from an exponential distribution with a minimum of 1 second, a maximum of 2 seconds, and a time constant (*τ*) of 1.2 seconds (**Fig. 4a**). Unless a behavior box malfunctioned and animals needed to be moved to a different one, experimenters made no interventions for these animals for over 100 days.

#### Sound Discrimination Task

We trained a cohort of rats (*N = 13*) to perform a sound discrimination task following fixation growth using the FixGrower curriculum (**Fig. 5**). The only differences in the curriculum occurred in stage 8 and are described below. Moreover, we provide detailed methods of how we added sounds into the task.

##### Sound Discrimination Stage 8– Penalty On

After reaching a target fixation duration of 1.45 seconds through the FixGrower fixation curriculum Stages 5–7, animals entered a modified version of Stage 8. Unlike in the original curriculum, this stage was not limited to a single session. Instead, animals remained in this stage until they met the following performance criteria in a single session: at least 250 consecutive trials with a violation rate below 20% and a hit rate above 85%. Once this criterion was met, they advanced to the sound discrimination phase.

Although the target fixation duration during training was 1.45 seconds, rats commonly maintained fixation beyond the go-cue, by 200–250 ms. This natural tendency to over-poke meant that animals were already capable of sustaining fixations close to the full task requirement of 1.65 seconds. To streamline training and eliminate the need for an additional session solely to reach this duration, we introduced the auditory stimuli concurrently with an increase in the fixation requirement to 1.7 seconds in the next stage. This strategy allowed us to reduce training time without compromising fixation stability during stimulus presentation.

##### Sound Discrimination Stage 9– Add Sounds

The goal of this stage was to introduce the two auditory tone stimuli (s_a_ and s_b_) during the fixation period while maintaining low violation rates. Animals continued to receive rewards via light-guidance following the go-cue, which corresponded to the correct choice based on the sound rule. Stimuli were introduced progressively—first by increasing duration, then by increasing volume.

Initially, tones were presented at 35% of maximum volume (20 dB), beginning at a duration of 30 ms and increasing within-session toward a target of 400 ms.Task timing was as follows: 250 ms pre-s_a_ delay→ 30 ms s_a_ → 940 ms pre-s_b_ delay → 30 ms s_b_ → 450 ms post-s_b_ delay, for a total fixation duration of 1.7 seconds. The inter-trial interval was sampled from a normal distribution with a mean of 1 second and a standard deviation of 0.5 seconds. Error choices triggered an additional 2-second timeout. Stimulus pairs were randomly drawn from two choice options and all tones were amplitude-matched to play at equal perceived volume (**Fig. 5b**):

1. **Left choice**: s_a_ = 3 kHz, s_b_ = 3 kHz
2. **Right choice**: s_a_ = 3 kHz, s_b_ = 12 kHz

On each trial, sound duration increased by 3 ms, while the delay between s_a_ and s_b_ decreased by 6 ms to maintain constant fixation time of 1.7 seconds. If the violation rate for a session exceeded 50%, sound growth was paused until the rate dropped below this threshold. If the violation rate remained below 40% for an entire session, the starting sound duration for the next session increased by 100 ms. This process repeated until the starting sound duration reached 400 ms, typically within four sessions. Final task timing was: 250 ms pre-s_a_ delay → 400 ms s_a_ → 200 ms pre-s_b_ delay → 400 ms s_b_ → 450 ms post-s_b_ delay (total fixation: 1.7 seconds).

After sounds played for their full duration, we gradually increased sound volume from 35% (20 dB) to 100% (80 dB). For the first 20 trials of each session, volume matched the previous session’s value. After 20 trials, volume increased by 13% (6.15 dB) and was held constant for the remainder of the session. Animals advanced past this stage once they achieved 100% volume with a hit rate above 85% and a violation rate below 30%. On average, animals completed this stage in 8.9 ± 1.75 days (min : 7,max : 13 days).

As a precaution, we introduced the sounds conservatively to avoid startling the animals. However, our data suggest this approach may have been overly cautious: animals tolerated the stimuli well, even during early exposures. Based on this observation, we recommend combining sound duration and volume increases in future training protocols, as animals appear capable of adapting to both manipulations concurrently.

##### Sound Discrimination Stage 10– Wean Off Light Guide

The next stage of training involved phasing out the light-guidance cue so that animals relied solely on the sound rule to make correct choices. Task timing was the same as at the end of Stage 9, with two modifications: the pre-s_b_ delay was reduced to 150 ms (total fixation: 1.65 seconds), and the penalty for error responses increased to a 4-second timeout in addition to the ITI.

In the first session of the stage we reduced the proportion of light-guided trials from 100% to 60%. At the end of each session, we computed accuracy on non-guided trials. If performance exceeded 70%, the light-guided trial fraction was reduced by 15% in the subsequent session. This process continued until, within a single session, the proportion of light-guided trials was less than 25%, accuracy on non-guided trials was at least 75%, and the violation rate remained below 30%. Once all criteria were met, animals advanced to the final stage. On average, this stage took 12.3 ± 4.1 days (min : 6,max : 21 days) to complete.

##### Sound Discrimination Stage 11– Sound Rule

In the final stage, animals were required to perform the task using only the auditory rule, with no light-guidance. This is the stage we analyze in **Fig. 5a-b**. All task parameters remained consistent with Stage 10 with the light guides turned off. Specifically, timing was: 250 ms pre-s_a_ delay → 400 ms s_a_ → 150 ms pre-s_b_ delay → 400 ms s_b_ → 450 ms post-s_b_ delay (total fixation: 1.65 seconds). Stimuli were randomly sampled from the set described earlier, requiring discrimination based on the frequency of s_b_ (**Fig. 5b**). The ITI remained normally distributed (mean = 1 second, SD = 0.5 second). Violations triggered a 2-second timeout in addition to the ITI, and error responses resulted in a 4-second timeout.

#### Mouse Fixation Task

We trained a cohort of mice (*N = 5*) to fixate using the FixGrower curriculum (Fig. 6). The only curriculum differences occurred in stage 8 and are described below.

##### Mouse Fixation Stage 8– Penalty On & Probe Violation Rates

After reaching a target fixation duration of 1.1 seconds through the FixGrower fixation curriculum Stages 5–7, mice entered a modified version of Stage 8. Unlike the original curriculum, the timeout duration following a violation was shortened to 100 ms and the ITI was sampled from a normal distribution with a mean of 2 seconds and a standard deviation of 0.5 seconds. Additionally, this stage was not limited to a single session and served as a benchmark for assessing violation rates (Fig. 6c-d). Mice remained in this stage until they met the following performance criteria in a single session: at least 150 trials with a violation rate below 20% and a hit rate above 85%. Once this threshold was met, animals completed the experiment.

#### Excluded Animals

Two animals from the fixation experiment were excluded from the final analyses, and their training progress is shown in (**Fig. S3**). Animal R044 (Legacy cohort) was excluded from all analyses related to the fixation growth and probe stages. This animal had substantial difficulty acquiring the center-poke behavior, requiring nine days and two behavior box changes before successfully executing a center-to-side motor sequence with a 10 ms hold. On the third day of successful trials, the behavior box malfunctioned, and the animal’s behavior failed to recover. Therefore, this animal is included only in analyses related to the side-poke training curriculum.

Animal R047 (FixGrower cohort) was excluded from analyses relating probe stage performance to training duration. This was the only animal in the dataset to trigger resetting logic in the fixation growth. R047 repeatedly engaged this mechanism during fixation growth, resulting in an unusually long training time (Days to Target = 22) despite exhibiting low probe-stage violation rates (9.0 ± 3%). As a result, this animal appeared as an outlier that broke the otherwise robust trend between training duration and performance observed in the FixGrower cohort. When included in the mixed-effects model, no significant interaction was detected between cohort and training time (*p* = 0.221). When excluded, a significant interaction emerged. This indicates that for animals that trigger the curriculum resetting logic, their training times may not be predictive of their final violation rate and alternative lines of evidence should be used to determine training potential.

### Simulated Fixation Growth

To assess how quickly animals in the Legacy cohort could have reached the 2 second fixation target under different behavioral assumptions, we simulated fixation growth using the Legacy curriculum algorithm under two conditions: (1) a **perfect** condition, in which no violations occurred, representing an upper bound on learning speed given each animal’s typical session length, and (2) a **recovery** condition, which incorporated each animal’s empirical violation patterns and session-level trial counts to validate the simulation against observed behavior.

#### Parameter Estimation

For each Legacy-trained animal, we estimated empirical session-level statistics using data from the fixation growth stages (Stages 5–7). These included the mean and variance of the number of trials per session, as well as warm-up and post–warm-up violation rates. Warm-up trials were defined as the first 20 trials of a session if the previous session’s final fixation duration exceeded 10 ms.

Violation rates were modeled using Beta distributions to capture their characteristic right-skew. Parameters for each animal’s warm-up and post–warm-up distributions were estimated via the method of moments. Specifically, for a given empirical mean (*μ*) and variance (*σ*^2^), Beta distribution shape parameters and *β* were derived using: 

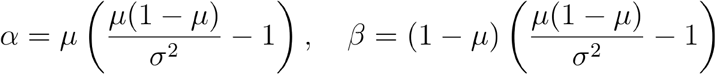

Session trial counts were sampled from a truncated normal distribution, with mean and variance set to empirical values, rounded to the nearest integer, and constrained to a minimum of one trial.

#### Simulation Procedure

For each simulated session, we first sampled the number of trials based on the estimated normal distribution of session lengths for that animal. In the perfect condition, both warm-up and post–warm-up violation rates were set to zero. In the recovery condition, violation rates were drawn from the animal’s corresponding Beta distributions for that session.

Each trial was simulated sequentially following the Legacy fixation growth logic. For each trial, we drew a Bernoulli sample using the appropriate violation probability (warm-up or post–warm-up) for the selected condition to determine whether the trial was a violation. For the first 20 trials of a session (warm-up phase), fixation duration increased linearly after each valid trial, with the step size calculated as a fixed increment designed to evenly span the gap between 10 ms and the ending fixation duration of the previous session. In the post–warm-up phase, the fixation duration increased by *max*(1 ms, 0.1%) each valid trial. Violation trials produced no change in the fixation duration.

The fixation duration at the end of each session was used as the warm-up target for the next session. Simulations continued until either the fixation duration exceeded 2 seconds or the simulation reached a maximum of 90 sessions. For each animal and condition, the simulation was repeated 30 times using a fixed random seed. Results were averaged across replicates to estimate the expected number of sessions required to reach the 2 second target in each condition.

### Trial Initiation Time

The trial initiation time is defined as the time from the final poke out from the center-port on the previous trial to the time of the first poke into the center-port on the current trial. To control for experimenter-imposed delays, we subtract off any incurred inter-trial-interval or penalty timeout that occurred on the previous trial.

### Statistical Analyses

Statistical analyses were performed using the SciPy and Statsmodels libraries in Python (3.10). Detailed statistical information is provided in the corresponding figure legends and Results sections. Unless otherwise specified, data are reported in-text as mean ± standard deviation (SD). Box plots show the first and third quartiles with the median indicated by a horizontal bar. Error bars and shaded regions in plots represent the standard error of the mean (SEM).

All statistical comparisons were conducted using non-parametric tests (e.g., *Mann–Whitney U* or *Wilcoxon signed*-*rank*) unless data were found to be normally distributed based on the *Shapiro–Wilk* test. When animals contributed multiple data points (e.g., session-level violation rates), mixed-effects linear models were used with animal ID included as a random intercept to account for within-subject variability. A significance threshold of *α* = 0.05 was used for all tests. Where applicable, *p-values* were adjusted for multiple comparisons using the *Holm*-*Bonferroni* method.

## Acknowledgments

We thank Klaus Osorio, Jovanna Teran, Andres Bustos and Emily Valance for technical assistance. We thank Grace Barnett and Jamus MacGuire for veterinary advice. We are grateful to all members of the Brody lab for their support, collegiality and feedback.

This manuscript was supported in part by the United States National Institutes of Health (NIH) under grants R01MH108358, R01MH138935, and 5U19NS132720. It is subject to the NIH Public Access Policy. Through acceptance of this federal funding, NIH has been given a right to make this manuscript publicly available in PubMed Central upon the Official Date of Publication, as defined by NIH. Additional support was provided by the Howard Hughes Medical Institute Investigator Program.

## Author Contributions

J.R.B. and J.A.C. conceived of and designed the study. J.R.B. and J.A.C. collected the data with help from

J.M.W. and C.D.K.. J.R.B. and J.A.C. analyzed the data. J.R.B and J.A.C. wrote the paper with feedback from C.D.B.. C.D.B. supervised the project and acquired funding.

## Data & Code Availability

All data will be published in a Zenodo database. All code for analyses will be published in a public repository on the Brody Lab Github.

## Competing Interests

The authors declare no competing interests.

## Supplemental Information

Figures S1-S7

Tables S1-S4

**Figure S1.**
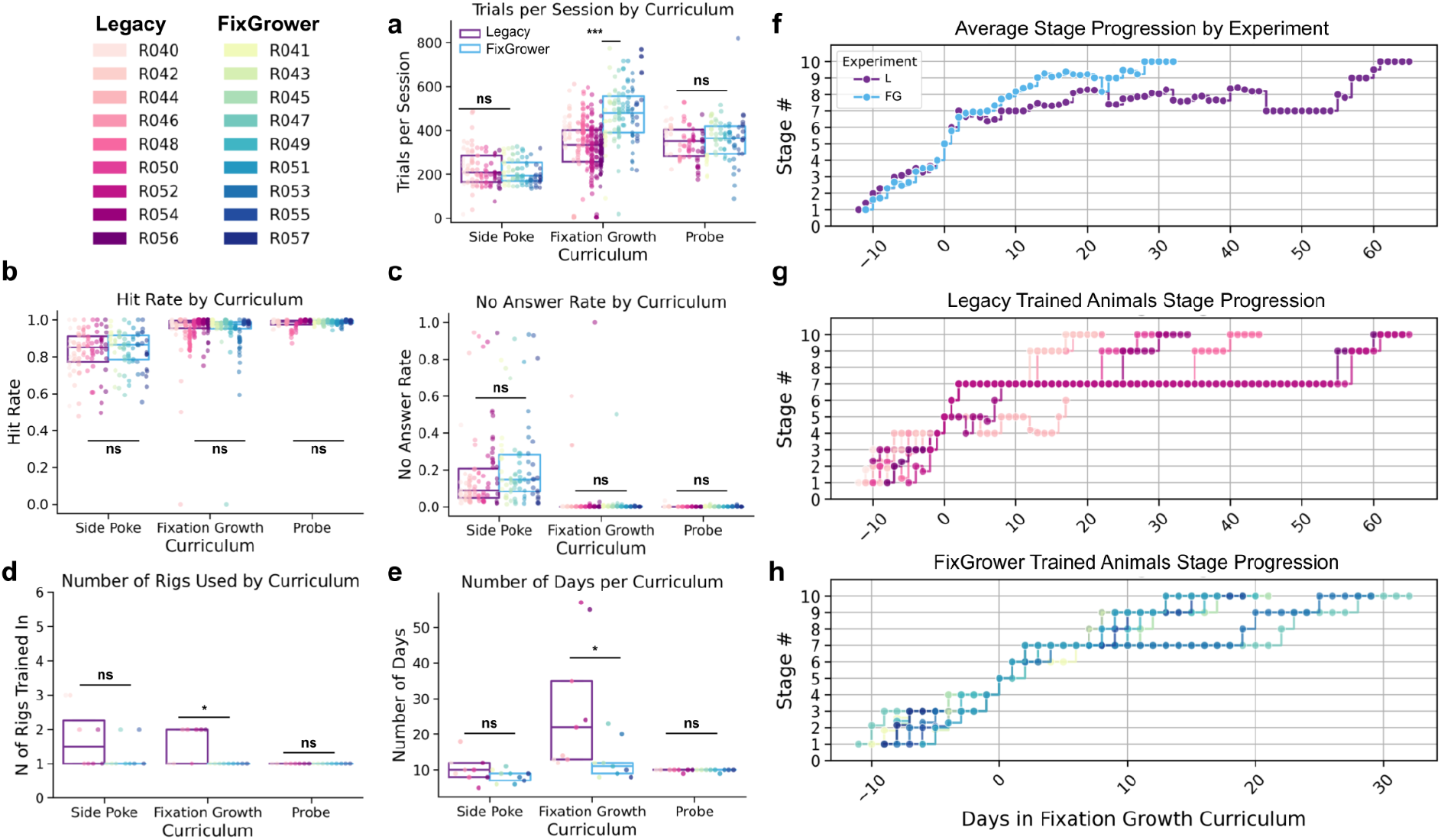
Comparison of task metrics across Curricula Phases: Side Poke, Fixation Growth, Probe. **(A)** Trials per session by curricula type compared by mixed linear model with a fixed effect for experiment and a random intercept for animal ID. Corrected for multiple tests with the Holm method. Experiment condition indicated by boxplot hue, individual animals indicated by dot color. Each dot indicates an animal’s performance in a session within a given stage type. **(B)** Same as **(A)** but for hit rate where a hit is defined as a valid trial with a correct side poke to the light guide. **(C)** Same as **(A)** but for no answer rate where no answer is defined as a valid trial with no side poke during the answer period (8 seconds post go-cue). **(D)** Number of behavior boxes (“rigs”) an animal trained in by stage type—see *Methods* for rig change rationale. Experiment condition indicated by box plot hue, individual animal indicated by dot color. Compared across experiment types by either *Mann–Whitney U* or *Welch’s t* depending on Shapiro normality and corrected for multiple tests by the Holm method. **(E)** Same as **(D)** but for the number of days spent in each stage type. **(F)** Average stage progression by experiment group aligned to the start of the fixation growth curriculum (Stage 5). **(G)** Individual stage progressions for all Legacy-trained animals aligned to the start of the fixation growth curriculum. **(H)** Same as **(G)** but for FixGrower-trained animals. *Statistics:* * *p < 0*.*05, ** p < 0*.*01, *** p < 0*.*001* and *ns* is *non-significant*.

**Figure S2.**
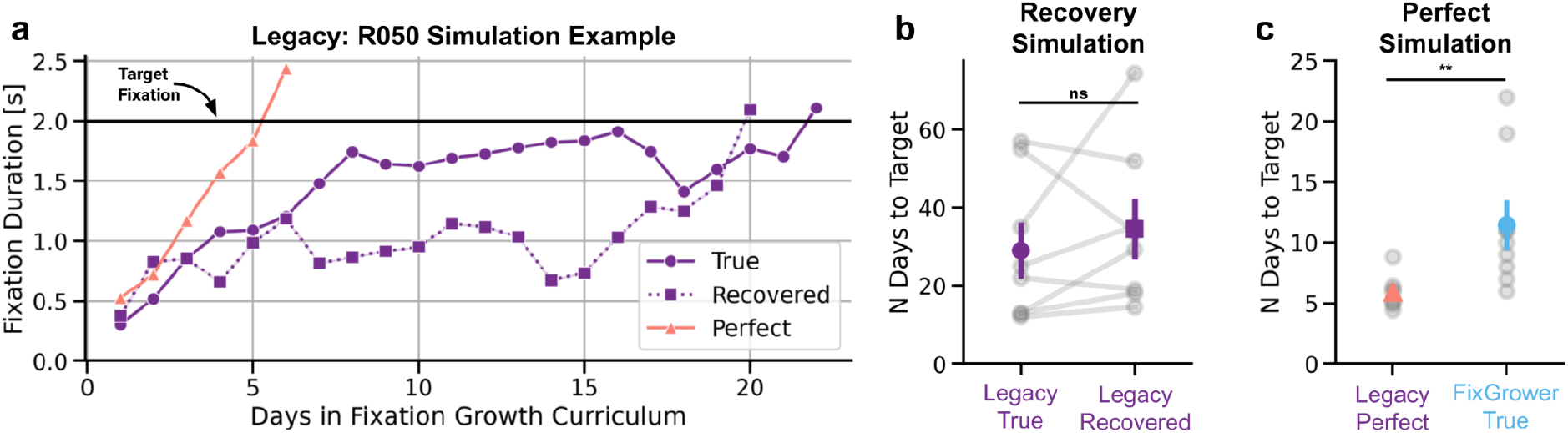
Simulations of Legacy-trained animals’ growth ceiling. **(A)** Example simulation for a single animal (R050), showing fixation duration over days for the true data (dark purple circles), recovery simulation (dashed purple squares), and perfect simulation (pink triangles). The black line marks the 2-second fixation target. See *Methods* for full simulation procedure. **(B)** Recovery simulation results for all Legacy-trained animals. Gray lines show individual animals (N = 9, *n*□_ee_d□ = 30) comparing true (circles) and recovered (squares) training durations. Purple markers indicate group means ± SEM. There was no significant difference in days to target between true and recovery conditions (*Wilcoxon signed-rank test, W* = 12.5, *p* = 0.547), supporting the validity of the simulation method. **(C)** Perfect simulation results for Legacy-trained animals (pink triangles) compared to true FixGrower-trained animals (light blue circles). Gray dots indicate individual animals; colored markers represent group means ± SEM. Under perfect conditions, Legacy-trained animals can reach the fixation target significantly faster than FixGrower-trained animals did, indicating they were not limited by an intrinsic growth-ceiling (*Mann–Whitney U, U* = 5.000, *p* = 0.003).

**Figure S3.**
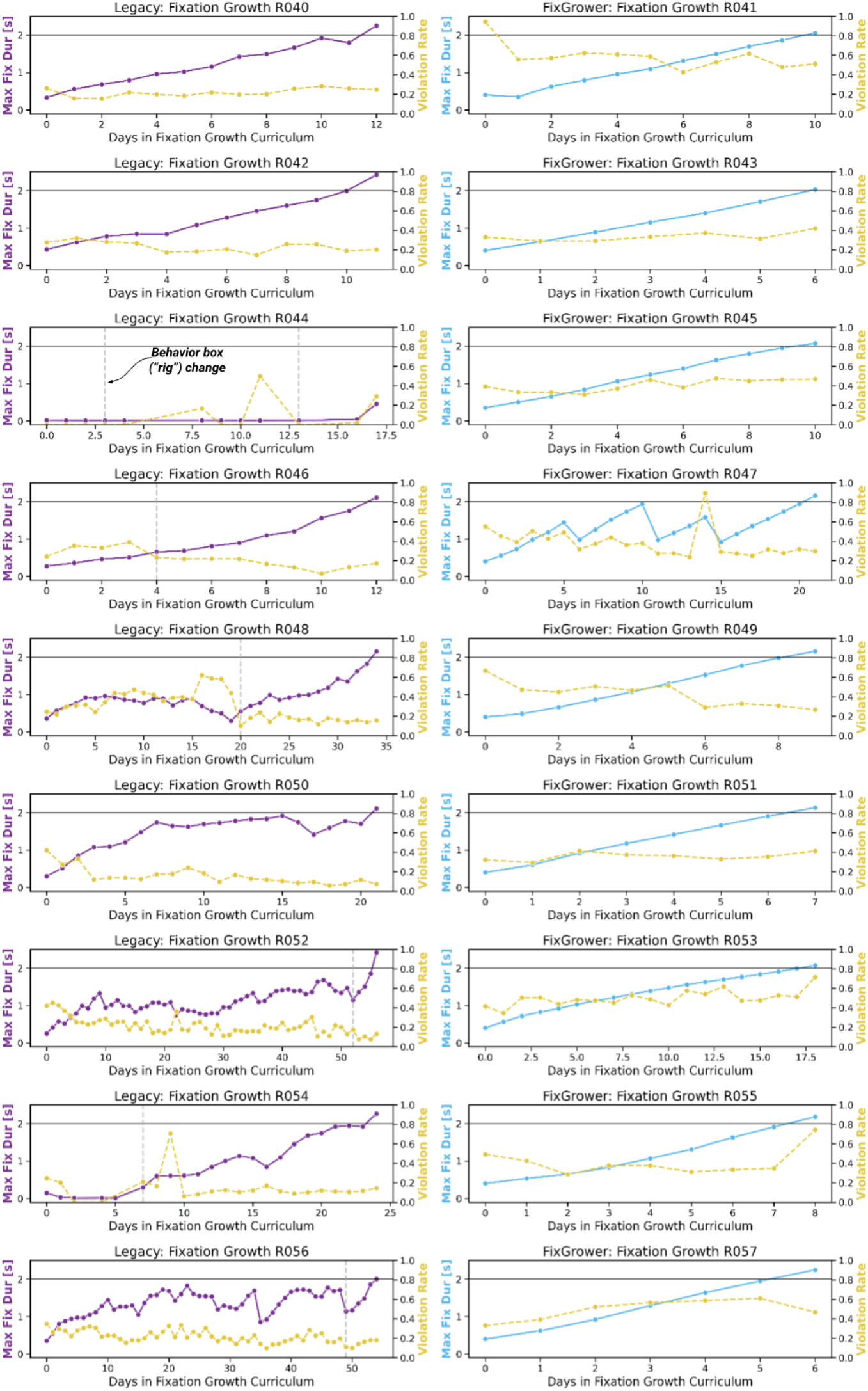
Raw fixation, violation and behavioral apparatus data. Data from sessions in fixation growth experiment for all Legacy and FixGrower-trained animals. The x-axis represents each day spent in the curriculum prior to reaching the 2-second fixation target. Left y-axis represents the maximum fixation duration reached in a given day, with the color (dark purple or light blue) indicating the experimental condition. Right y-axis indicates the violation rate for the given day. Gray dashed vertical lines indicate days in which a behavioral apparatus (“rig”) was switched. Animal Legacy:R044 was dropped from analysis of growth timing and fixation performance due to equipment error and discontinued training. Animal FixGrower:R047 was an outlier in Figure 3.3 analysis.

**Figure S4.**
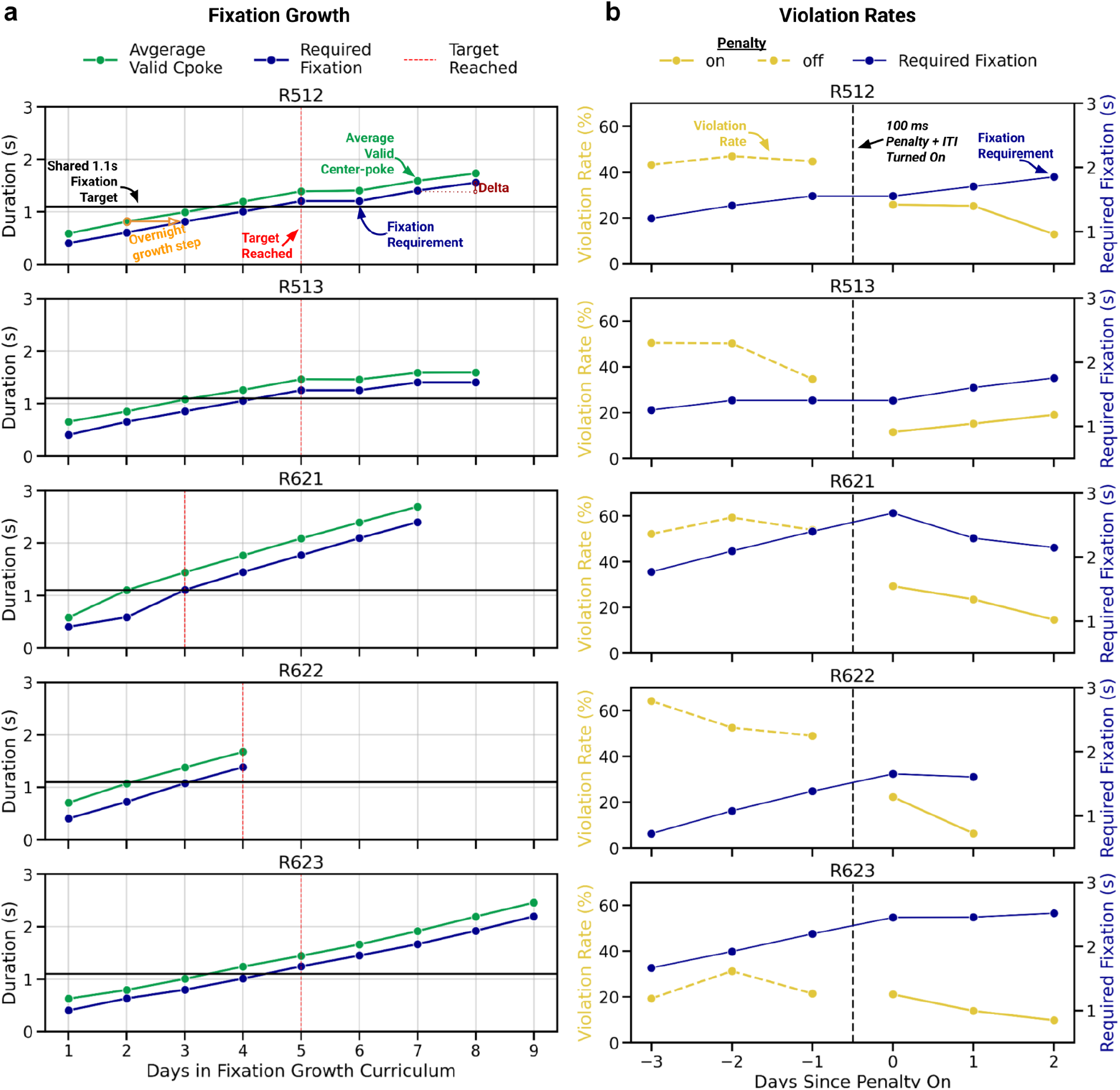
Extended data from mouse fixation experiment. **(A)** Fixation duration growth curves for individual mice across days in FixGrower. Green lines represent the average duration of valid center-pokes per day that was used to determine the next day’s requirement (blue). Red dotted vertical lines mark the session in which each mouse first reached the shared 1.1-second fixation target (horizontal black line). The top panel (R512) includes annotations illustrating the curriculum’s overnight stepwise growth (orange), and the session-to-session delta (maroon) used to assess growth rate. **(B)** Violation rate trajectories for each mouse before and after the onset of a 100 ms penalty and inter-trial interval (ITI). Yellow lines denote daily violation rates, with dashed lines representing pre-penalty sessions and solid lines post-penalty. The black dashed vertical line indicates the session when the penalty was introduced. Required fixation duration is overlaid in blue. Number of post-penalty sessions represents the number of days it took animals to reach a violation rate below 20% and perform at least 150 trials.

**Figure S5.**
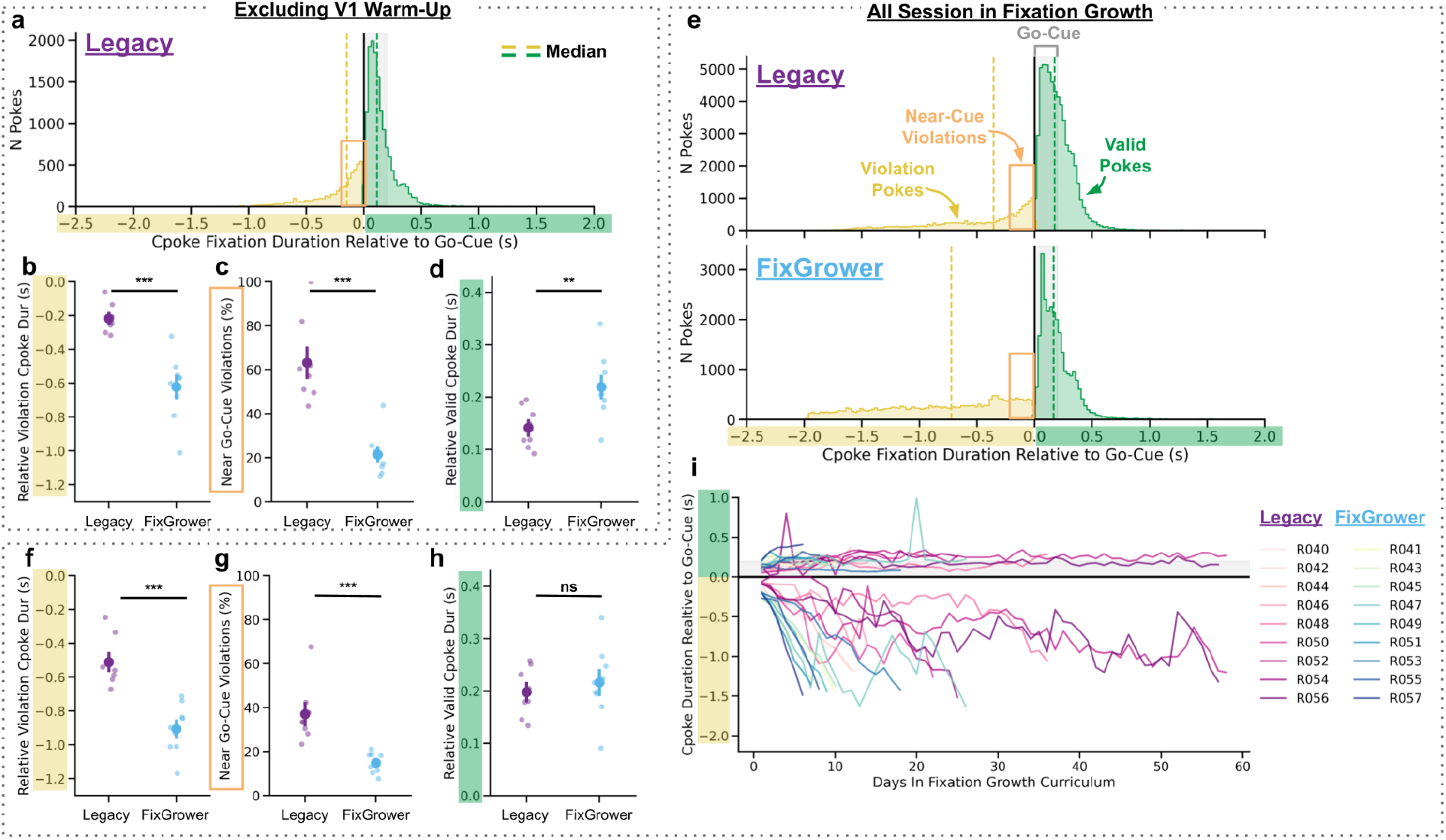
Additional analysis accounting for warm-up exclusion and full fixation growth duration. **(A)** Distribution of center-poke fixation durations relative to the go-cue for Legacy-trained animals after excluding the initial 20-trial “warm-up” period. Valid pokes are shown in green, violations in yellow. Dashed lines indicate medians. The orange box indicates the near-cue period (200 ms preceding the go-cue), used for quantification in **(B, G). (B–D)** Same metrics as in Figure 3.9B–D, computed after excluding the 20-trial warm-up for Legacy-trained animals. FixGrower animal data did not change from Figure 3.9. Data is only from the first 7 days of fixation growth. **(B)** Mean violation poke times relative to the go-cue; **(C)** percent of violations occurring within the near-cue window; and **(D)** valid poke durations relative to the go-cue. Statistical tests do not differ when warm-up trials are excluded. **(E)** Poke duration distributions for valid and violation trials across the full fixation growth period for both Legacy (top) and FixGrower (bottom) animals, which is an extension from the first 7 days shown in Figure 3.9. **(F–H)** Same as **(B–D)**, but calculated from all sessions and trials during fixation growth. **(I)** Center-poke durations for average valid and violation pokes relative to the go-cue across fixation growth days for individual animals in both cohorts. Each line represents a single animal. *Statistics:* All scatter/point plots show mean ± SEM with individual animal data points overlaid. * *p < 0*.*05, ** p < 0*.*01, *** p < 0*.*001* and *ns* is *non-significant*

**Figure S6.**
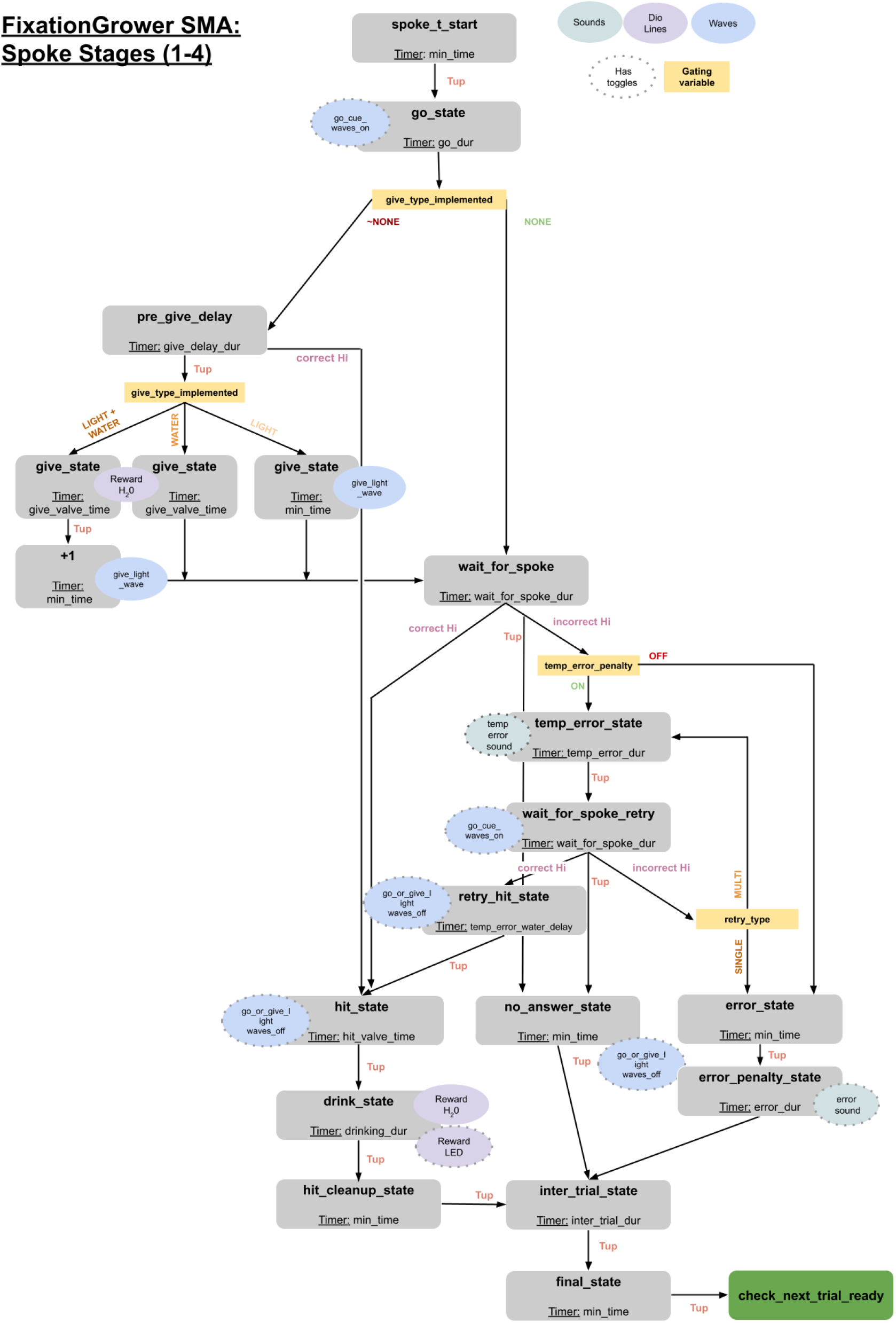
Visualization of the state machine logic written in Bcontrol for the side-poking curriculum stages. Arrows indicate state machine transitions and are labeled by the condition they require. ‘Tup’ refers to the timer for a state running to completion.

**Figure S7.**
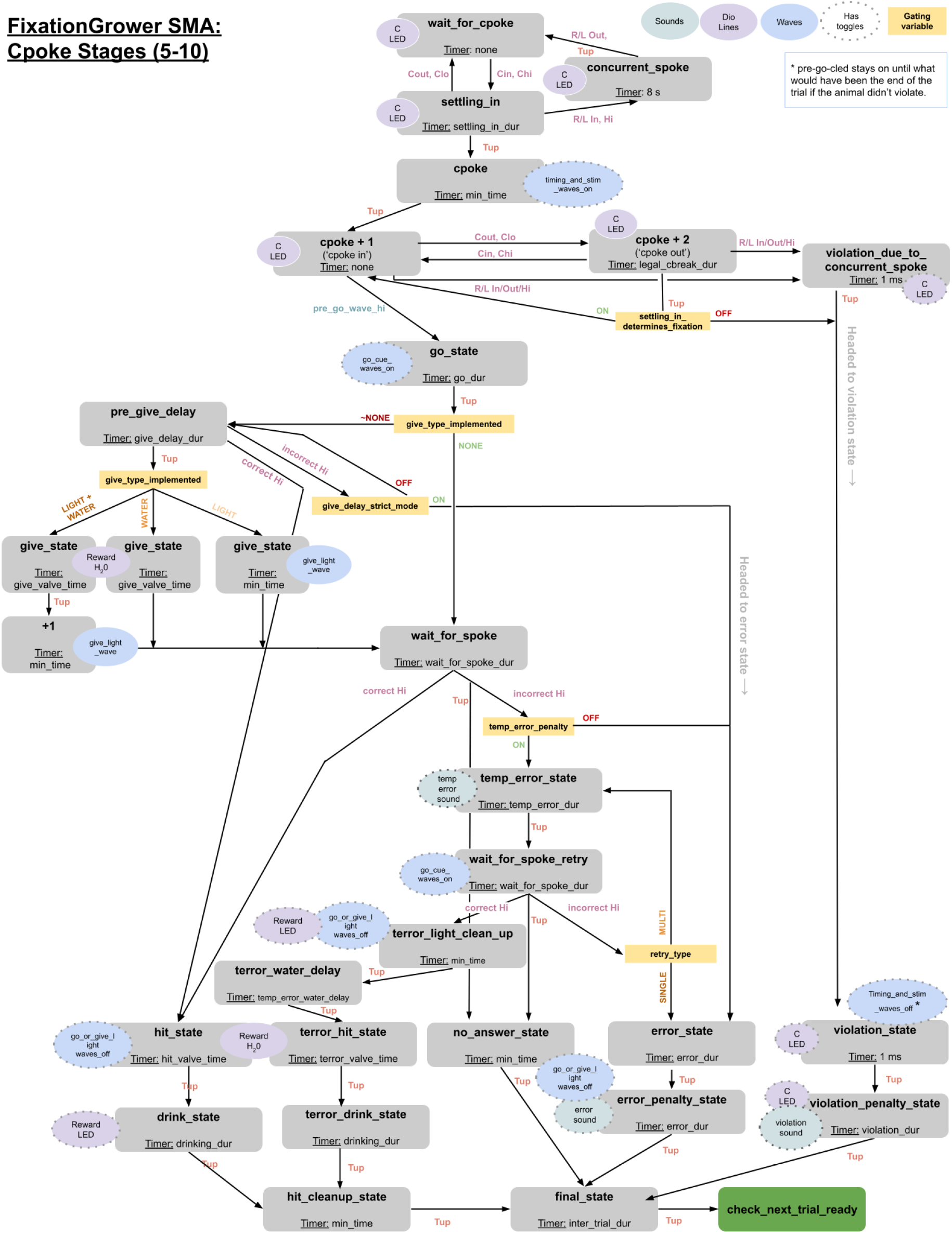
Visualization of the state machine logic written in Bcontrol for the center-poking stages. Fixation Experiment stages 5–11). Arrows indicate state machine transitions and are labeled by the condition they require. ‘Tup’ refers to the timer for a state running to completion.

**Table S1.** Overall description of training stages. Similarities and differences are noted for each stage. Red indicates the stages of the fixation growth curriculum with stages that differ between conditions.

| Stage | Description | Shared | Legacy Only | FixGrower Only |
| --- | --- | --- | --- | --- |
| 1 | Learn L Spoke | All Logic |  |  |
| 2 | Learn R Spoke | All Logic |  |  |
| 3 | Alternate L/R Spoke Blocks | All Logic |  |  |
| 4 | L/R Spoke Blocks → Random | All Logic |  |  |
| 5 | Center Fixation to Side Blocks with Growth | Trial structure, blocks, stage completion logic | Fix. start @ 10ms, grow by valid trial, violation penalty on | Fix. Start @ 400 ms, Grow overnight |
| 6 | Center Fixation to Side Blocks → Random with Growth | Trial structure, blocks, stage completion logic | Fixation warm up, grow by valid trial, violation penalty on | Grow overnight |
| 7 | Center Fixation to Side → Random with Growth | Trial structure, stage completion logic | Fixation warm up, grow by valid trial, violation penalty on | Grow overnight |
| 8 | FG Violation Penalty On | None- FG only |  | Violation penalty on |
| 9 | Probe: Stable Fix. Delay | All Logic |  |  |
| 10 | Probe: Random Fix. Delay | All Logic |  |  |

**Table S2.** Summary of variable parameters across side poke learning stages. The go-cue and side reward LED are present in all stages. Under “Sides,” AB denotes anti-bias. Under “Water,” values in parentheses indicate mouse reward volumes. Under ITI “N(X,Y)” indicates sampling from a normal distribution with mean X and standard deviation Y. Under “Completion,” ‘t’ refers to trials, and EOD (End of Day) indicates that stage transitions occur only at the end of the session and are implemented in the following session.

| Stage | Side(s) | Water | Retry | ITI | Completion |
| --- | --- | --- | --- | --- | --- |
| 1 | L | Seed 18 (3) μL,<br>Reward 42 (10) μL | Y | N(40 s, 5 s) | > 40 t, NA < 15%<br>within session |
| 2 | R | Seed 18 (3) μL<br>Reward 42 (10) μL | Y | N(40 s, 5 s) | > 40 t, NA < 15%<br>within session |
| 3 | L/R 15 t blocks | Reward 45 (6) μL | N | N(25 s, 3 s) | @EOD > 90 t,<br>Hit > 87%, NA < 20%, |
| 4 | L/R 15 t blocks →<br>rand w/ AB @ 70 t | Reward 30 (3) μL | N | N(15 s, 3 s) | @EOD > 175 t,<br>Hit > 80%, NA < 20% |

**Table S3.** Overview of training parameters that differ between the two fixation learning stages. The two training curricula differed in their initial fixation values, their growth algorithms and their violation penalties.

|  | **<u>Legacy</u>** | **<u>FixGrower</u>** |
| --- | --- | --- |
| **Initial Fixation Value** | Every Day: 0.01 s | Day 1: 0.4 s<br>Day > 1: previous session's average valid cpoke duration |
| **Growth Algorithm** | By valid trial:<br>max(1 ms, 0.1%) following 20 trial warm up towards yesterday's fix. maximum | By session:<br>To the average valid cpoke duration of the most recent session |
| **Violation Penalty** | 2 second timeout followed by ITI | No penalty or ITI until stage 8, then 2 second timeout followed by ITI |

**Table S4.** Detailed overview of stage-wise variable changes shared by both Legacy and FixGrower curricula. Stage 8 is excluded, as it applies only to FixGrower-trained animals. “AB” denotes anti-bias; “t” indicates trials; “EOD” (End of Day) signifies that stage transitions occur only overnight.

| Stage | Side(s) | Completion |
| --- | --- | --- |
| 5 | L/R 15 t blocks | @EOD 200 t, Hit > 85%, NA < 10% |
| 6 | L/R 15 t blocks → rand w/ AB @ 70 t | @EOD 250 t, Hit > 85%, NA < 10% |
| 7 | rand w/ AB | @EOD 200 t, Hit > 85%, NA < 10%,<br>final fixation dur > target |

## Notes

### Competing Interest Statement

The authors have declared no competing interest.

